# The ventral CA2 region of the hippocampus and its differential contributions to social memory and social aggression

**DOI:** 10.1101/2024.06.07.597964

**Authors:** Lara M. Boyle, Wanhui Sheng, Felix Leroy, Rhea Sahai, Sarah Irfan, Heon-Jin Lee, Andres Villegas, W.Scott Young, Steven A. Siegelbaum

**Author notes:** These authors contributed equally to this work (Lara M. Boyle, Wanhui Sheng). Corresponding and senior author: Steven A. Siegelbaum.

## Abstract

Although it is well-known that the hippocampus mediates declarative memory (the repository of information of people, places, things and events) and influences behavior, the differential contributions of the dorsal and ventral hippocampus to specific forms of memory and behavior remain largely unknown. Studies to date show that the dorsal hippocampal CA1 region is important for cognitive and spatial tasks whereas the ventral CA1 region is associated with affective or emotional processing. Whether other regions and forms of hippocampal-dependent memory and behavior show a similar distinction remains unclear. Here we examine how social memory and related social behaviors are encoded across the dorso-ventral axis of the hippocampus. Although recent studies show that the dorsal hippocampal CA2 region is required for social memory and acts to promote social aggression, the behavioral role of ventral CA2 remains unknown. Indeed, whether a defined CA2 region extends throughout ventral hippocampus is controversial. Here, we report that a molecularly, anatomically and electrophysiologically defined CA2 region extends to the extreme ventral pole of hippocampus, with both similarities and important differences in its projection patterns and synaptic impact compared to dorsal CA2. Of particular importance, we find that ventral CA2 is not required for social memory but is critical for promoting social aggression. These results support the view that the ventral region of hippocampus is more generally tuned for emotionally-related behaviors compared to the cognitively-tuned dorsal hippocampus.

## Introduction

One of the key goals of hippocampal research is to elucidate how the different subregions of this extended structure interact to regulate different forms of declarative memory. In addition to the distinct regions of dentate gyrus, CA3, CA2, and CA1 along the transverse axis of the hippocampus, as initially defined by Lorenté de Nò^1^, the dorsal and ventral regions along the longitudinal axis of the hippocampus differ in their gene expression, connectivity, and behavioral role^2^. A comprehensive RNA sequencing study of the hippocampus found that the transcriptional distance between dorsal and ventral CA1, CA3, and DG are approximately as large as those seen between the different transverse subfields^3^.

Functionally, the dorsal region of the hippocampus has been linked to contextual memory and spatial navigation^4,5^. In contrast, disruptions in ventral hippocampus function cause changes in anxiety-like behavior and emotional processing^6–9^. These different behavioral roles are reflected in the marked differences in the extrahippocampal projections from dorsal and ventral CA1. Thus, whereas dorsal CA1 (dCA1) projects mainly to subiculum, entorhinal cortex and lateral septum, ventral CA1 (vCA1) uniquely projects to prefrontal cortex, nucleus accumbens, amygdala and hypothalamus. Although dorsal and ventral CA3 do not differ markedly in their extrahippocampal projections, they do show differences in their functional role in memory and behavior^10–12^, potentially via topographic projection patterns through CA1.

The distinction between the behavioral roles of dorsal and ventral hippocampus is of particular interest for social memory, which involves the storage and recall of information about other individuals of the same species. The importance of the hippocampus for social memory has been evident since the studies by Brenda Milner and her colleagues of patient HM^13^, and appears differentially regulated along the dorso-ventral axis of the hippocampus. Optogenetic silencing of dCA1^14^ or chemogenetic silencing of dCA3^15^ or dorsal dentate gyrus^16^ does not impair social novelty recognition memory (SNRM), the behavioral discrimination of a novel from familiar conspecific. In contrast, optogenetic silencing of vCA1 and chemogenetic silencing of vCA3 suppresses SNRM recall and encoding, respectively.

Although the results on CA1 and CA3 suggest a selective role for ventral hippocampus in social memory, a number of studies have now found that the CA2 region is critically important for the encoding, consolidation and recall of social memory^17–23^. In addition, CA2 directly regulates social aggression through a trisynaptic circuit to the lateral septum and hypothalamus. However, to date, all published studies of the behavioral role of CA2 have focused on its dorsal portion (dCA2). Despite descriptions of ventral CA2 (vCA2) by Lorente de Nó^1^ and others^24–26^, the existence of vCA2 has been a matter of controversy^27^.

CA2 pyramidal neurons were first identified by Lorenté de Nò^1^ based on their distinct morphology and anatomical location between CA1 and CA3. More recent work has focused on the unique electrophysiological properties of dCA2 pyramidal neurons relative to neighboring dCA1 and dCA3 neurons^28–30^ as well as the distinct profile of gene expression in CA2 relative to neighboring CA1 and CA3, including CA2-selective expression of RGS14, PCP4, and Amigo2. Of particular note, CA2 is highly enriched in receptors for the social neuropeptides arginine vasopressin (AVP)^31^ and oxytocin (OXT)^32^, where they contribute to regulating social behaviors. Thus, general deletion of the arginine vasopressin receptor 1b (Avpr1b)^33^ or the oxytocin receptor^32^ impairs social memory, as does the conditional deletion of both receptors when restricted to CA2^23^. In addition, general deletion of Avpr1b inhibits social aggression^33^ and this phenotype can be rescued by selective viral expression of Avpr1b in dCA2 and nearby neurons^34^, indicating that AVP acting on Avpr1b expressed in dCA2 neurons is necessary to promote normal levels of aggression.

The results concerning the identification of CA2 pyramidal neurons in more ventral regions of hippocampus have been less clear cut and have relied mostly on gene expression patterns. Although certain dorsal CA2 markers, including PCP4 and Avpr1b, have been found to extend into ventral hippocampus, other markers do not. In addition, markers that are reliably co-expressed in dorsal CA2 are less reliably expressed in ventral CA2^27,31,35^. Thus, the full extent to which CA2 extends along the dorsoventral axis is unclear. Based on in-situ hybridization data from the Allen Brain Atlas, CA2 was suggested to be restricted to the dorsal two-thirds of the hippocampus and to be absent from the ventral one-third of the hippocampus^36^. However, because of the curved shape of the hippocampus, it can be difficult to identify the anatomical location of CA2 relative to CA1 and CA3 along the transverse axis of the hippocampus in the coronal and sagittal slices presented in the Allan Brain Atlas. A more recent meta-analysis of in situ hybridization and single cell mRNA expression data from the Allen Brain Atlas and DropViz databases suggested that the expression of at least a subset of CA2 markers may extend further towards the ventral pole^37^, although the pattern of molecular expression in ventral CA2 neurons may be distinct from their dorsal counterparts^37^. At present, little is known as to the electrophysiological properties and synaptic connectivity of putative CA2 neurons in the most ventral region of hippocampus, including whether such neurons share the characteristic properties of dorsal CA2. Importantly, the behavioral role of ventral CA2, and whether it participates in social behaviors as defined for dorsal CA2, remains unexplored.

Here, we performed a multi-disciplinary investigation of the molecular properties, electrophysiological properties, synaptic connectivity, and behavioral role of ventral CA2 using Avpr1b-Cre^26^ and PCP4-Cre (RBRC05662, developed by Toshitada Takemori and provided by RIKEN BRC) mouse lines. We found that the expression of Avpr1b and PCP4, and their respective Cre-driven expression, extends along the entire dorsoventral axis of the hippocampus, while other CA2 molecular markers are more restricted to the dorsal and intermediate segments of the hippocampus. Moreover, whereas many CA2 markers defined to date are co-expressed in dCA2 pyramidal neurons, there is considerable molecular heterogeneity in vCA2. While pharmacogenetic inhibition of pyramidal neurons in dCA2 in both mouse lines results in loss of social novelty recognition memory, inhibition of pyramidal neurons in vCA2 does not alter social recognition behavior in either mouse line. In contrast, inhibition of vCA2 reduces social aggression, similar to the effect of silencing dCA2. These different behavioral roles are mirrored in complementary patterns of synaptic connectivity from dCA2 and vCA2 to their vCA1 intrahippocampal and dorsal lateral septum extrahippocampal targets. Thus, ventral CA2 is a unique region within the hippocampus with distinct molecular expression, connectivity, and a functional role that is separate from the canonical role in social memory behavior associated with dorsal CA2.

## Results

### The boundaries of CA2 extend along the entire dorsoventral hippocampus

Given the differing reports on ventral CA2 noted above, a question arises: how should CA2 be defined in the hippocampus? Here we have defined CA2 pyramidal neurons based on the following features: 1) anatomical location between CA1 and CA3, spanning the end of the mossy fibers from dentate gyrus, 2) lack of thorny excrescences that characterize CA3 pyramidal neurons, 3) restricted or enriched expression of genes and/or proteins not expressed in CA1 or CA3, including RGS14, STEP, and/or Avpr1b, and 4) distinct electrophysiological properties that differ from CA1 or CA3^29,30^.

We first examined the expression of tdTomato in the *Avprb1-Cre* x Ai14 tdTomato reporter line along the entire dorsoventral axis of the hippocampus using either horizontal brain slices (Fig, 1) or transverse slices of dissected hippocampus (Fig. 2). In both preparations, the transverse axis is well-preserved, facilitating the identification of CA2. We found that tdTomato was expressed along the entire extent of the dorsoventral axis at a location along the transverse axis corresponding to the expected location of CA2 between CA3 and CA1(Fig. 1). We quantified the density of Avpr1b-expressing tdTomato+ neurons in the anatomically defined CA2 region as a fraction of DAPI-positive cells (Figure 2). There was a significantly smaller fraction of tdTomato+ cells in the CA2 region of ventral hippocampus (38.0±5.4% of DAPI-labeled cells) compared to dorsal hippocampus (72.4±8.8% of cells; two-sample unpaired t-test t=8.953, df=13, p<0.0001), confirming previous qualitative observations^26^.

**Figure 1.**
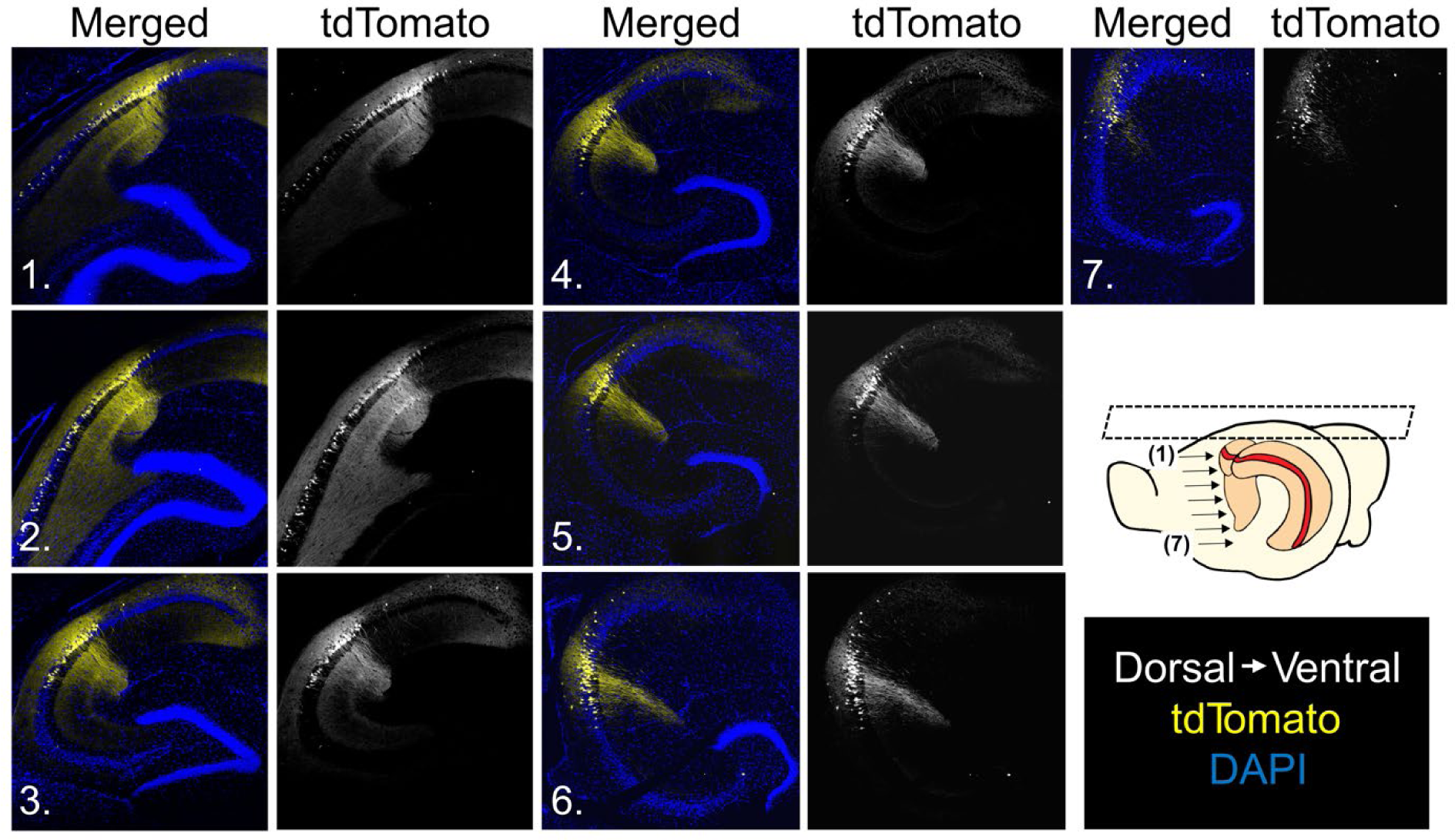
Horizontal slices from Avpr1b-Cre x Ai14 mice reveal tdTomato expression along the entire dorsoventral axis. tdTomato (Avpr1b) expression is present along the entire axis from the most dorsal (1) to most ventral (7) slices. Numbers on images correspond to arrows in inset.

**Figure 2.**
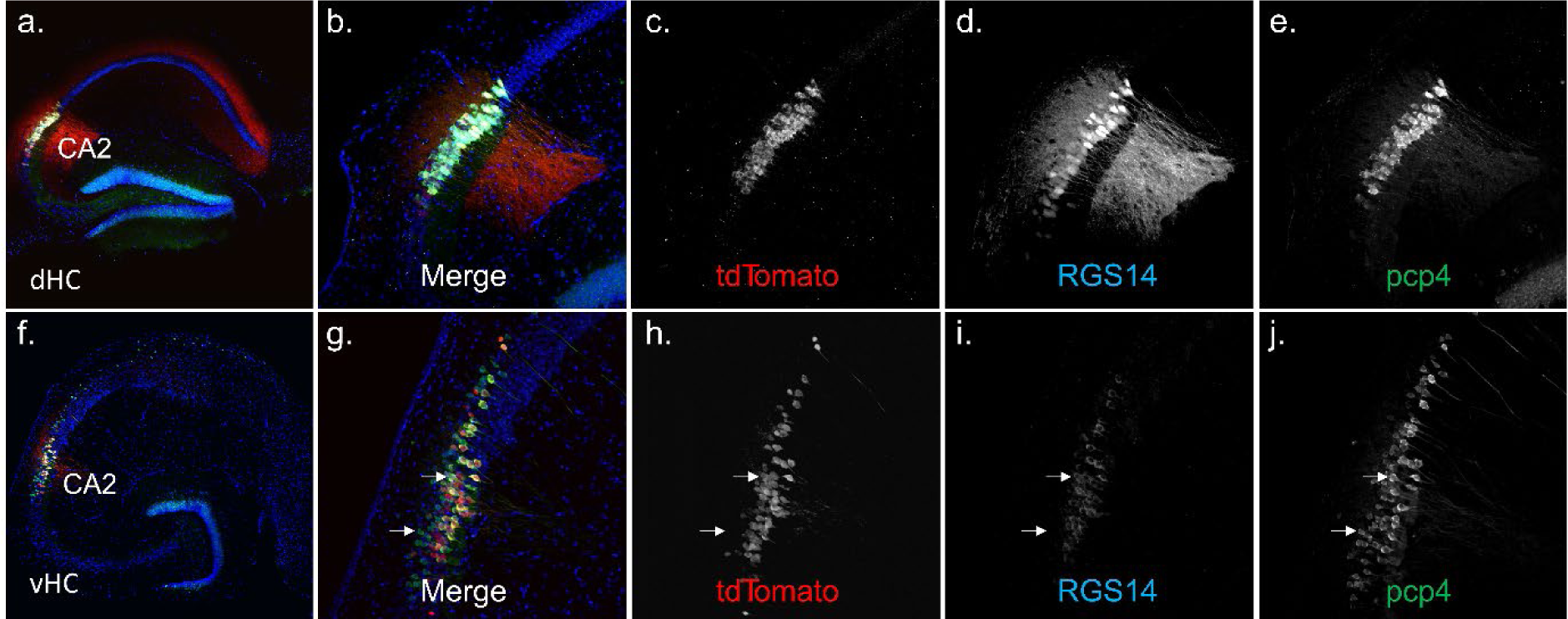
Distinct patterns of co-expression of prototypical CA2 markers and Avpr1b-Cre mediated expression in dorsal compared to ventral hippocampus. **a-e)** Dorsal hippocampus transverse slices (n=8) from Avpr1b-Cre x Ai14 mice (n=4) labeled with RGS14, PCP4, and DAPI. **f-j)** Ventral hippocampus transverse slices (n=7) from Avpr1b-Cre x Ai14 mice (n=3) labeled with RGS14, PCP4, and DAPI. Top white arrow indicates an example cell expressing two markers plus tdTomato (triple-labeled). Bottom white arrow indicates an example cell that expresses PCP4 but not tdTomato (Avpr1b) or RGS14.

To determine whether Avpr1b-Cre driven expression was continuous along the dorsoventral axis, we employed the brain clearing technique iDisco and light sheet microscopy to observe the expression pattern along the entire dorsoventral axis simultaneously. We found that Avpr1b was indeed expressed in a continuous band from the most dorsal to the most ventral portions of the hippocampus (Suppl. Fig. 1).

We next examined whether tdTomato expression overlaps with *Avpr1b* gene expression in the adult *Avpr1b-Cre* mouse by performing in-situ hybridization in *Avpr1b-Cre* x *Ai14* mouse brains (n=2), using probes against *Avpr1b* and *tdTomato* mRNAs in coronal sections. We found that Avpr1b and tdTomato transcripts were fully colocalized, demonstrating that Cre-driven expression accurately reports the site of Avpr1b expression in the ventral hippocampus in the adult mouse (Suppl. Fig 2), similar to results in dorsal hippocampus^26^.

We next examined whether *Avpr1b-Cre* mediated expression was confined to neurons expressing other CA2 pyramidal cell markers along the dorsal-ventral axis. In dorsal hippocampus there is near complete overlap between all molecular markers for CA2, including *Avpr1b-Cre* mediated expression^26^. Although there is evidence for increased heterogeneity in expression of certain CA2 markers in ventral hippocampus, the extent to which the different dorsal CA2 markers are co-expressed in ventral CA2 neurons has not been examined. We thus performed immunofluorescent staining and imaging of several markers of dCA2 that have been shown to be expressed in the ventral hippocampus, including RGS14, PCP4 and STEP, along the entire dorsoventral axis in transverse slices from the dissected hippocampus of two cohorts of *Avpr1b-Cre* x *Ai14* mice. In one cohort, we co-stained slices for RGS14, PCP4, and the nuclear marker DAPI (Fig. 2). In another cohort, we co-stained slices for STEP, PCP4, and DAPI (Suppl. Fig. 3a). Further, since PCP4 labels the dentate gyrus and mossy fiber tract, we confirmed that vCA2, like dCA2, has an anatomical localization centered at the distal end of the mossy fibers (Suppl. Fig. 4). A consistent region was defined within dorsal and ventral CA2 to standardize comparison of positive cell counts across these regions (see Methods). This area (Suppl. Fig. 4) was selected because >50% of DAPI-stained neurons within this boundary express the putative CA2 marker PCP4. We quantified the total number of cells in the region that were co-labeled by at least one putative CA2 marker and DAPI (Figure 2).

Similar to results with the *Avpr1b-Cre* x *tdTomato* reporter line, the percent of DAPI-expressing cells in the anatomically defined CA2 region in ventral hippocampus that expressed at least one of the examined CA2 markers (60.9 ± 3.9%) was significantly less than observed for dorsal hippocampus (72.8 ± 8.7%, two-sample unpaired t-test t=3.333, df=13, p=0.0054). Moreover, whereas the percent of tdTomato-expressing cells in dorsal CA2 (72.4 ± 8.8%) was nearly identical to the percent of dCA2 cells expressing any of the CA2 markers (72.8 ± 8.7%), there were significantly fewer cells in ventral CA2 expressing tdTomato (38.0 ± 5.4%) than the percent of cells expressing any of these markers (60.9±3.8%), indicating the presence in vCA2 of a subpopulation of tdTomato-negative (ie, Avpr1b-Cre-negative) cells that expressed at least one or more of the other CA2 markers (Suppl. Fig. 5).

To further understand this difference, we quantified the number of cells that co-expressed each of the combinations of these markers in e*Avpr1b-Cre* x *tdTomato* mice (Figure 3). In dCA2, 98.2 ± 1.4% of cells that expressed one of these prototypical markers co-expressed the two other CA2 markers examined (tdTomato, PCP4, and either RGS14 or STEP). In vCA2, PCP4 was the most widely expressed marker, present in 60% of DAPI-positive cells. Moreover, PCP4 was co-expressed in all cells expressing one or more of the three other CA2 markers examined.

**Figure 3.**
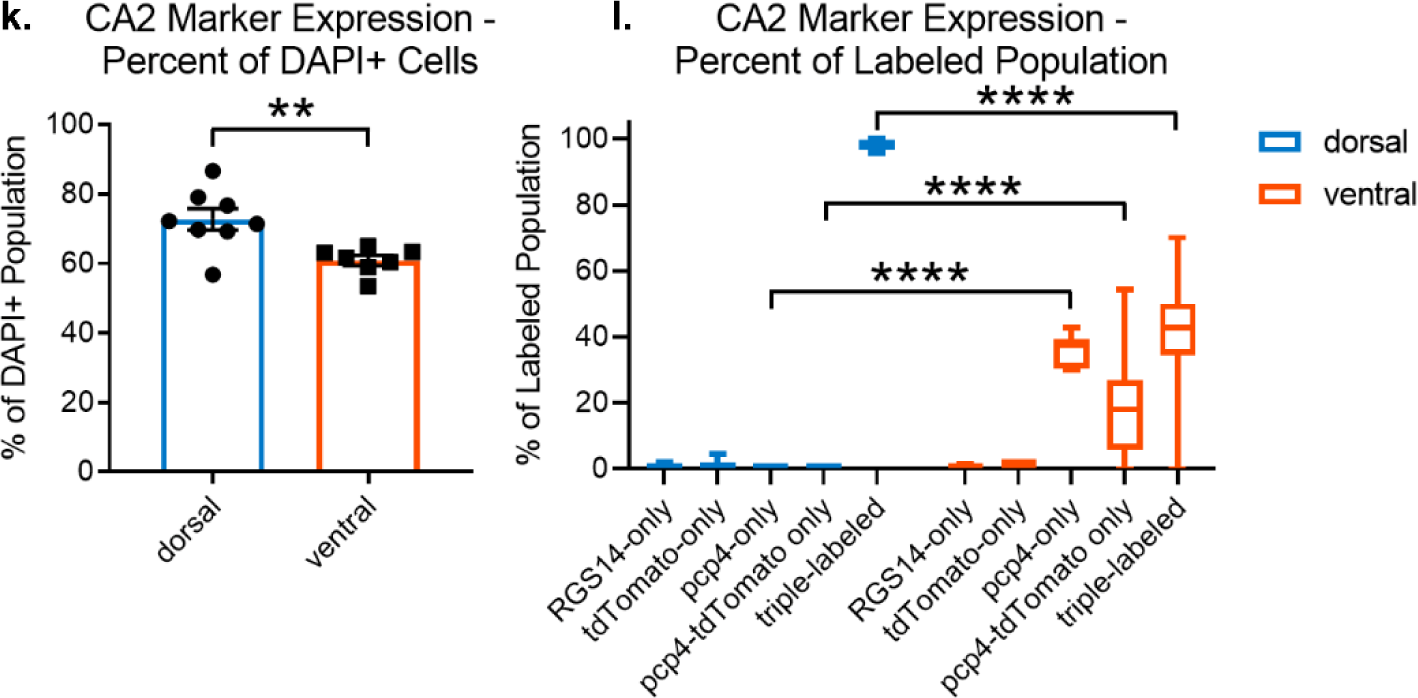
a) Quantification of percent of cells expressing DAPI within CA2 that express at least one of the prototypical CA2 markers (tdTomato/Avpr1b, PCP4, or RGS14). Significantly fewer cells in the ventral hippocampus express at least one of these markers. **b)** Percent of labeled cells that express each of the prototypical CA2 markers, or combinations therein. In the dorsal hippocampus, 98.2 ± 1.4% of cells express three prototypical CA2 markers, while virtually none (0.0 ± 0.0%) expressed only PCP4. In the ventral hippocampus, significantly fewer cells express three markers (41.5 ± 19.9%) with greater variance, and significantly more cells only express PCP4 (36.6 ± 6.0%) compared to dCA2. **p<0.01, ****p<0.0001

Furthermore, in 36.6 ± 6.0% of pyramidal cells in the vCA2 region, PCP4 was the only CA2 marker that these neurons expressed. In contrast, in dorsal CA2 there were few if any cells that only expressed PCP4. The PCP4-only cells appeared to be primarily in the deep pyramidal cell layer (defined as cells located closer to the stratum oriens). Thus, the majority of vCA2 neurons expressed PCP4, while a significantly smaller fraction expressed tdTomato and RGS14/STEP (Suppl. Fig. 3). Altogether, these data reveal that pyramidal neurons expressing dCA2 markers extend throughout the ventral region of the hippocampus, furthering previous results^37^.

### Electrophysiological and anatomical properties of vCA2 pyramidal neurons resemble those of dCA2

In addition to the patterns of gene expression that distinguish dorsal CA2 from its CA1 and CA3 neighbors, dorsal CA2 pyramidal neurons also have a number of characteristic electrophysiological membrane properties that distinguish them from CA1 and CA3 neurons^29,38^. Given the molecular heterogeneity we observed between dCA2 and vCA2 neurons, we wondered whether the intrinsic electrophysiological properties of these populations also differed. We thus performed whole cell patch clamp recordings from pyramidal neurons located in the anatomical CA2 region in acute slices from ventral hippocampus. When we compared our vCA2 recordings with results from prior studies from our lab on dCA2, we found that the two groups of cells exhibited nearly identical intrinsic electrophysiological properties, including input resistance, membrane capacitance and amplitude of voltage sag during membrane hyperpolarization, a hallmark of the hyperpolarization-activated cation current I_h_ (Suppl. Fig. 6). Of further note, the electrophysiological properties of ventral CA2 neurons were clearly distinct from those we observed in ventral CA1 (Suppl. Fig. 6). Thus, compared to ventral CA1, neurons in ventral CA2 had a lower voltage sag in response to a hyperpolarizing current step, indicating a lower I_h_, a lower input resistance, a higher rheobase, and a higher membrane capacitance, consistent with the larger soma of CA2 compared to CA1 pyramidal neurons.

Dorsal CA2 neurons also differ from dCA1 in their relative synaptic responses to their two major excitatory inputs: the direct perforant path inputs from entorhinal cortex that target the most distal apical dendrites in stratum lacunosum moleculare (SLM) and the Schaffer collateral inputs from CA3 neurons that target the more proximal apical dendrites in stratum radiatum (SR). Patch clamp recordings from dCA2 have shown that the depolarizing postsynaptic potential (PSP) evoked by stimulation of the direct EC inputs is much greater than the PSP evoked by activation of the Schaffer collateral inputs, the inverse of what is observed in dCA1^29^. To examine the synaptic responses of vCA2 neurons, we performed whole cell recordings from transverse ventral hippocampal slices from *Avpr1b-Cre x Ai14* reporter mice, targeting the CA2 region of the slice based on tdTomato fluorescence (although we did not target specifically the tdTomato-positive neurons), while activating inputs with a stimulating electrode in SLM or SR (Figure 4).

**Figure 4.**
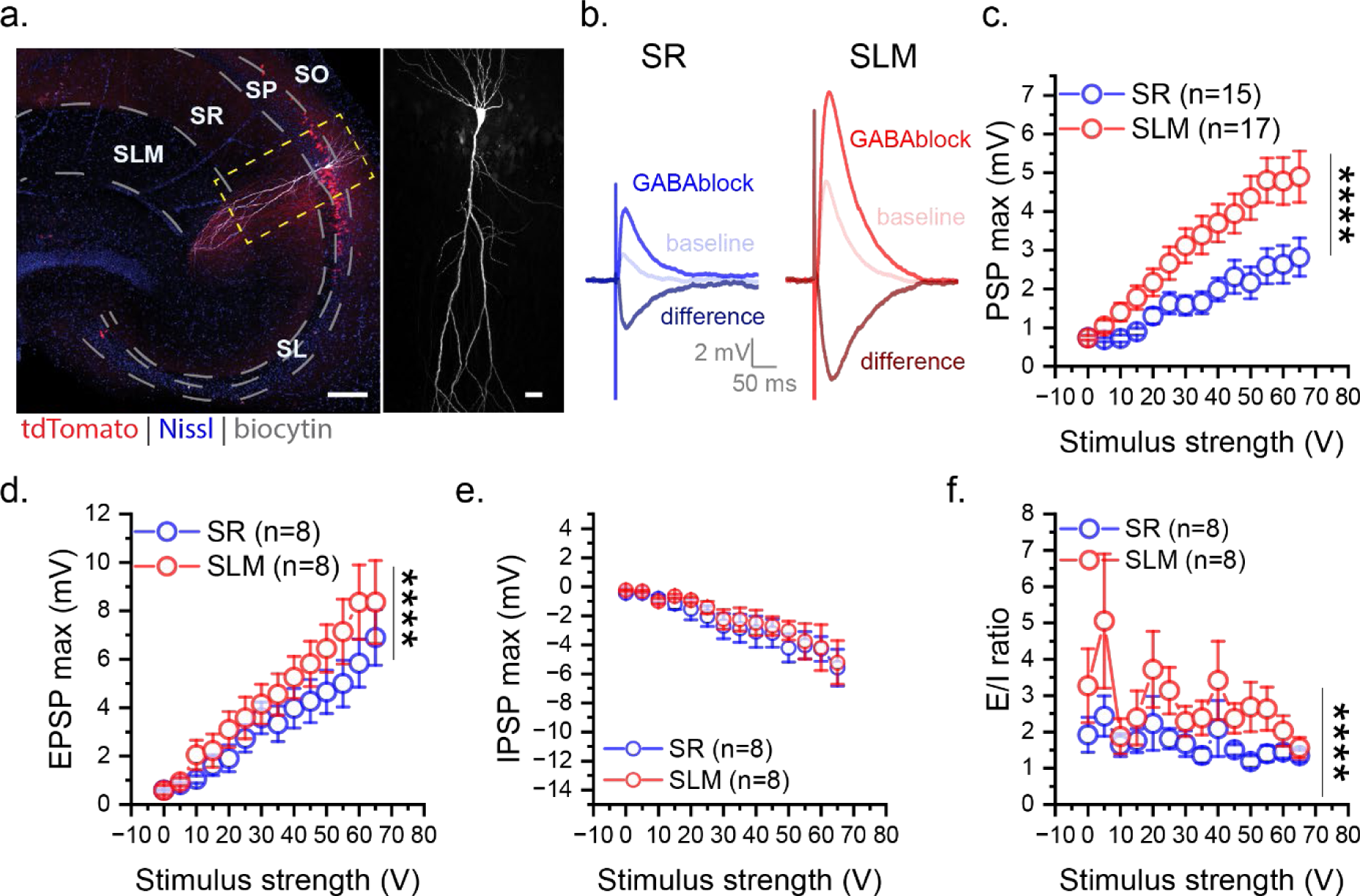
vCA2 pyramidal neurons receive stronger excitatory inputs from EC than from CA3. **a)** A transverse ventral hippocampal slice with a single biocytin-filled CA2 pyramidal neuron (white) from an AVPR1b-Cre x Ai14 mouse (left). Higher magnification view showing the morphological features of the same cell (right). Scale bars: 200 µm, left; 30 µm, right. **b)** Sample traces of the compound PSPs (baseline), the EPSPs recorded after applying GABA antagonists (GABAblock) and the IPSPs obtained by subtracting PSPs from the paired EPSPs (difference). EPSPs elicited with stimulating electrodes placed in SR (blue traces) and SLM (red traces). **c-e)** The amplitude of the PSP (Two way ANOVA F(1, 420)=81.04, p<0.0001, SR: n_mice_=9, SLM: n_mice_=10) and EPSC (e, Two way ANOVA F(1, 196)=16.73, p<0.0001, SR: n_mice_=5, SLM: n_mice_=6) evoked from EC (SLM) was significantly larger than that evoked from CA3 (SR) in vCA2 pyramidal neurons; same amplitude of IPSC (Two way ANOVA F(1, 196)=1.973, p=0.1617, SR: n_mice_=5, SLM: n_mice_=6). **f)** The EPSP/IPSP amplitude ratio from the populations of cells shown in (e-f) (Two way ANOVA F(1, 196)=19.09, p<0.0001, SR: n_mice_=5, SLM: n_mice_=6). ****p<0.0001

Similar to results in dCA2 (Chevaleyre and Siegelbaum, 2010), we found that vCA2 pyramidal neurons generated a much larger depolarizing PSP in response to electrical stimulation in the SLM compared to the PSP in response to stimulation in SR (Fig. 4b,c). As the PSP is the net voltage response generated by the excitatory postsynaptic potential (EPSP) in response to activation of glutamatergic inputs and the inhibitory postsynaptic potential (IPSP) in response to activation of GABAergic inputs by the electrical stimulation, we dissected out the EPSP and inferred IPSP by applying antagonists of GABA_A_ receptors (SR-95531) and GABA_B_ receptors (CGP-35348) to block inhibitory synaptic responses. Similar to results in dCA2 (Chevaleyre and Siegelbaum, 2010), the EPSP elicited by SLM stimulation was greater than that elicited by SR stimulation (Fig. 4b,d). We then calculated the inferred IPSP by subtracting the EPSP from the net PSP, and found similar inhibitory responses to electrical stimulation in SLM and SR (Fig 4b,e). As a result the EPSP to IPSP ratio was greater with SLM stimulation compared to SR stimulation (Fig. 4f).

Lorenté de Nò distinguished CA2 neurons from CA3 neurons based on the absence or presence, respectively, of large dendritic spines (thorny excrescences) in stratum lucidum, which are the site of the mossy fiber synapses from dentate gyrus granule cells^1^. In contrast, most CA2 neurons typically lack such large spines (although cf ^39^). We therefore assessed the dendritic morphology of vCA2 neurons by performing patch-clamp recordings with biocytin in the patch pipette from neurons in ventral hippocampal slices in Avpr1b-Cre x tdTomato mice. Post-hoc examination of biocytin-filled tdTomato-positive cells following electrophysiological recordings revealed that their dendrites were lacking in detectable thorny excrescences, similar to findings in dCA2 (Suppl. Fig. 7). To confirm the differences between vCA2 and vCA3 morphology, we obtained patch-clamp recordings using biocytin-filled pipettes from neurons in the ventral CA3 region and examined their dendritic processes. In contrast to tdTomato-expressing neurons, biocytin-filled tdTomato-negative neurons in ventral CA3 reliably exhibited large spines in stratum lucidum characteristic of thorny excrescences (Suppl. Fig. 8).

### vCA2 pyramidal neurons project to distinct areas within vCA1 and lateral septum compared to dCA2

Next we examined whether dorsal and ventral CA2 project to similar or distinct brain regions. Dorsal CA2 sends dense projections to dCA1 and dorsal lateral septum, with somewhat weaker projections to ventral CA1, with the latter two outputs implicated in promoting social aggression^40^ and enabling social memory^20^, respectively. To date, the outputs of genetically identified ventral CA2 neurons remain unknown.

To compare dCA2 and vCA2 projections, we performed anterograde tracing by injecting *Avpr1b-Cre* mice with Cre-dependent viruses expressing distinct fluorescent marker proteins (see Methods) in dCA2 (GFP or YFP-expressing virus, n=4) and vCA2 (RFP-expressing virus, n=4). We then imaged transverse, horizontal, and coronal sections to reveal the vCA2 and dCA2 connectivity pattern within and outside the hippocampus (Figures 5,6).

**Figure 5.**
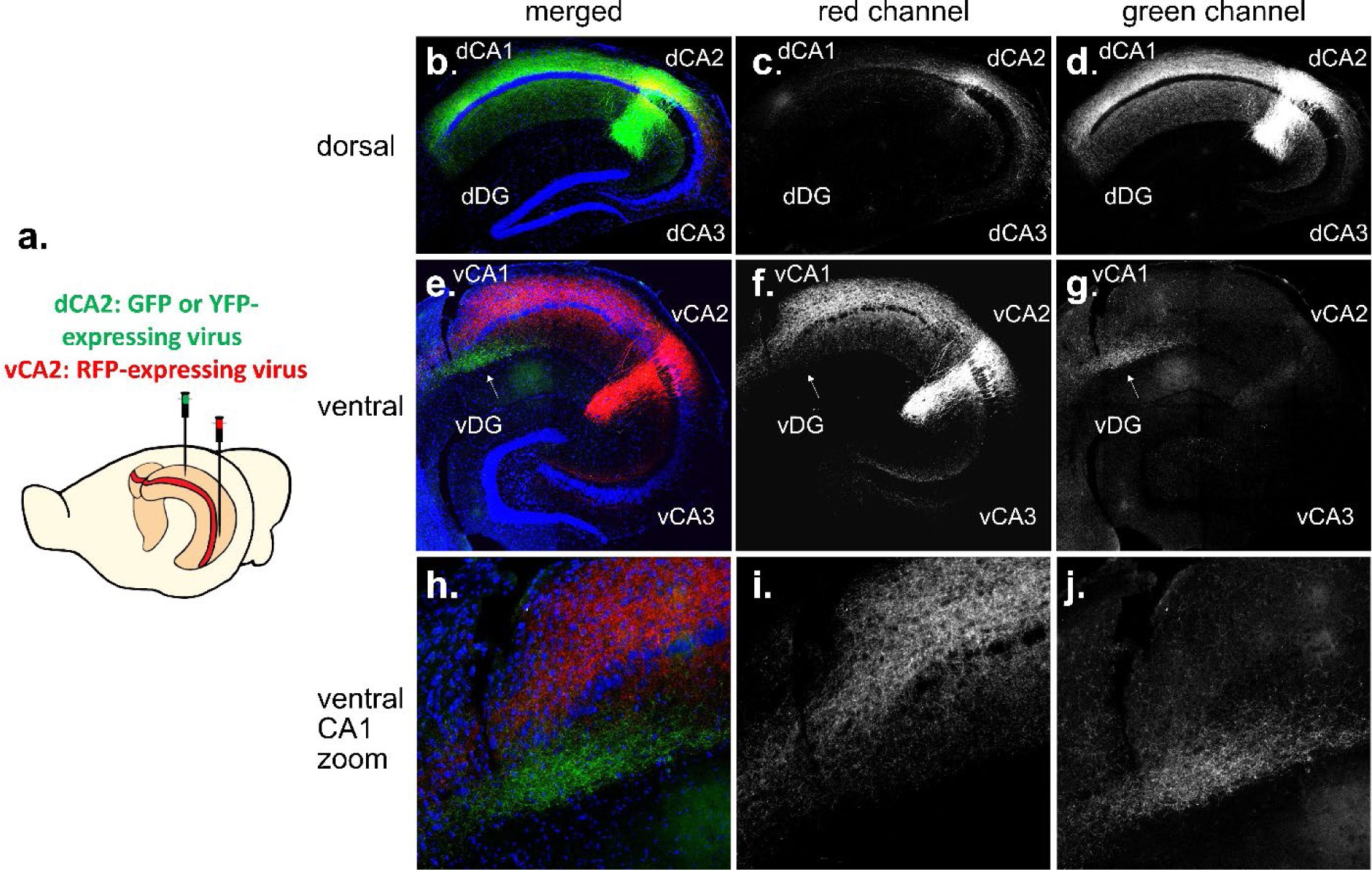
Dual injection in dCA2 and vCA2 reveals a topographic projection pattern along the hippocampus dorsoventral axis. **a)** Experimental scheme showing Cre-dependent GFP or YFP-expressing virus injected in dCA2 and Cre-dependent RFP-expressing virus injected in vCA2 in Avpr1b-Cre mouse. Transverse slices in dorsal hippocampus **(b-d)** and ventral hippocampus **(e-j)**. b,e) Composite tile images of injection site in dCA2 **(b)** and vCA2 **(e). c,f)** Red-channel showing vCA2 projections to dHC **(c)** and vHC **(f)**. Green-channel showing dCA2 projections to dHC **(d)** and vHC **(g)**. **h)** Zoomed image of vCA1 from composite image in **e)** showing separate projection pattern of dorsal and ventral CA2 to ventral CA1. **i)** red channel (vCA2) projection to vCA1 is predominantly located in stratum orients. **j)** green channel (dCA2) projection to vCA1 is predominantly located in stratum radiatum. White arrow indicates magnified vCA1 region in h-j.

**Figure 6.**
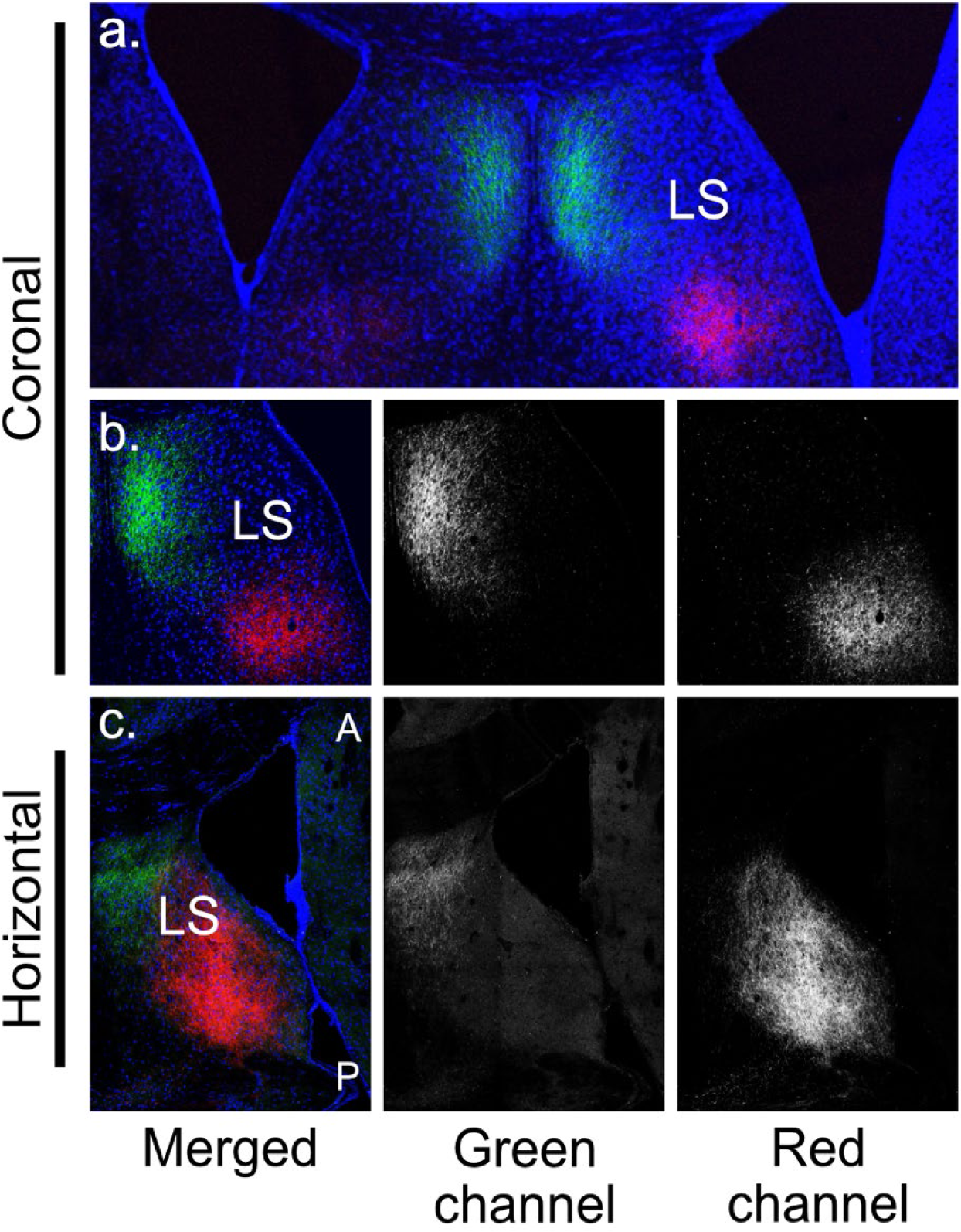
Dual-injection experiment reveals topographic projections from CA2 to the lateral septum. **a)** 5x composite image of a coronal slice from an Avpr1b-Cre injected with Cre-dependent AAV to express EYFP in dCA2 and Cre-dependent AAV to express mCherry in vCA2. Images show projections from dCA2 (green) to anterior dorsal lateral septum and from vCA2 (red) to posterior ventral lateral septum. **b, left)** 20x merged image of projections from dCA2 (green) and vCA2 (red) to lateral septum. **b, center)** Green channel (dCA2 projections) only. **b, right)** Red channel (vCA2 projections) only. **c, left)** 10x horizontal tile from Avpr1b-Cre mouse showing projections from dCA2 (green) to anterior dorsal lateral septum and projections from vCA2 (red) to posterior ventral lateral septum. White A indicates anterior side; white P indicates posterior side. **c, center)** Green channel (dCA2 projections) only. **c, right)** Red channel (vCA2 projections) only.

We observed a topographic pattern of intrahippocampal projections along the CA2 dorsoventral axis (Fig. 5), with dCA2 projecting primarily to dCA1 and vCA2 projecting primarily to vCA1.

Interestingly, the dCA2 projections to vCA1 target a region complementary to the vCA1 region targeted by vCA2. Thus, dCA2 projects to a limited region of vCA1 bordering the subiculum (distal vCA1 or vCA1a). In contrast, vCA2 projects to a wider region of vCA1, including the region of vCA1 bordering vCA2 (proximal vCA1 or vCA1c) and the middle region of vCA1 (vCA1b), with projections to vCA1a, the region targeted by dCA2, being relatively sparse. The projections from dCA2 and vCA2 to vCA1 also target complementary layers of vCA1. Thus, whereas dCA2 axons are largely confined to stratum radiatum (as previously described, Meira 2018), vCA2 axons largely confined to stratum oriens, the same region in which the projections from dCA2 to dCA1 are largely confined (Fig. 5g, j), as previously described^20^.

Similar to dCA2, we find that the primary extrahippocampal output from vCA2 targets the lateral septum. As described for other hippocampus inputs to lateral septum^41^, the CA2 septal projection is topographic, with dCA2 projecting to dorsal anterior lateral septum, and vCA2 projecting to a more ventral posterior region of the lateral septum, as previously described using non-genetic based tracer approaches^37^ (Fig. 6).

Next we performed electrophysiological recordings to determine the synaptic strength of the vCA2 outputs to vCA1. We injected *Avpr1b-Cre* animals with Cre-dependent AAV to express ChR2 in vCA2 pyramidal neurons and compared the optogenetically-evoked postsynaptic potentials recorded in pyramidal neurons from vCA1a with the responses recorded in vCA1b/c, the site of densest projections from vCA2 (Fig. 7). We found that light evoked a measurable postsynaptic depolarization in a significantly greater fraction of vCA1b/c pyramidal neurons compared to vCA1a pyramidal neurons (Fig 7b; Chi-square, z=2.338, p=0.019). In addition, optogenetic stimulation evoked a significantly larger peak depolarization and larger positive time-integral voltage response in vCA1b/c neurons compared to vCA1a (Fig. 7d-e, Two-way ANOVA vCA1a versus vCA1b/c; PSP max: F(1, 174)=21.99, p<0.0001; PSP positive integral: F(1, 174) = 9.833, p=0.002).

**Figure 7.**
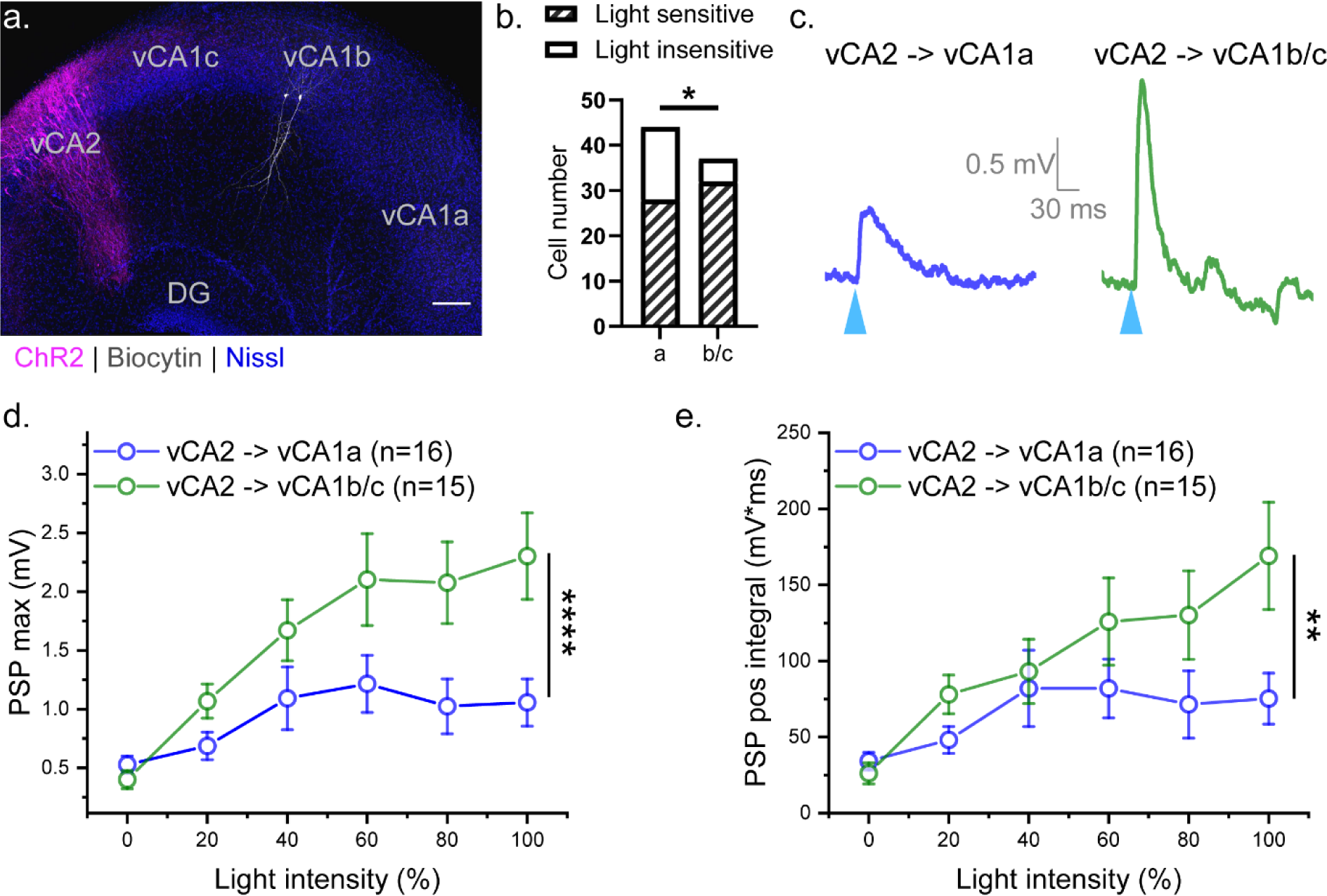
Optogenetic activation of vCA2 evokes larger PSP in pyramidal neurons in vCA1b/c compared to vCA1a. **a)** Avpr1b-Cre+/- mice were injected with ChR2 in vCA2. Light-evoked PSP was measured in post-synaptic pyramidal neurons located in vCA1a or vCA1b/c. White neurons in vCA1b show biocytin staining following patch clamp recordings with biocytin in electrode. Scale bar, 100 µm. **b)** Proportion of neurons in vCA1a and vCA1b/c with measurable light-evoked synaptic responses in slices expressing ChR2 in CA2 (Chi-square, z=2.338, p=0.019). Cells recorded from voltage clamp and current clamp were pooled. **c)** Example traces of light-evoked PSP in a vCA1a and vCA1b/c neuron. **d)** Maximum PSP amplitude (Two-way ANOVA vCA1a vs. vCA1b/c: F(1, 174)=21.99, p<0.0001) and **e)** integral of PSP (Two-way ANOVA vCA1a vs. vCA1b/c: F(1, 174) = 9.833, p=0.002) were significantly greater in vCA1b/c compared to vCA1a. **p<0.01, ****p<0.0001

As the PSP represents the net effect of the direct monosynaptic excitatory postsynaptic response to vCA2 activation and any feedforward disynaptic inhibitory response, we performed voltage clamp recordings of the excitatory and inhibitory postsynaptic currents (EPSCs and IPSCs) in vCA1 neurons in response to optogenetic activation of vCA2 neurons. For comparison, we also measured the voltage-clamped EPSCs and IPSCs from dCA1 neurons in response to optogenetic activation of dCA2. Optogenetic activation of dCA2 and vCA2 produced similar peak EPSCs in dCA1 and vCA1, respectively, with no significant difference in the responses (Suppl. Fig. 11a). In contrast, optogenetic activation of vCA2 elicited a significantly larger IPSC in vCA1 neurons, compared to the IPSC recorded in dCA1 in response to optogenetic activation of dCA2 (Suppl. Fig. 11b). As a result, the inhibitory/excitatory synaptic current ratio (I/E ratio) is significantly larger in vCA1 in response to activation of vCA2 compared to the ratio in dCA1 in response to activation of dCA2 (Suppl. Fig. 11d).

Thus, Avpr1b-Cre expressing cells in the ventral hippocampus meet many of the morphological, molecular, electrophysiological and synaptic properties of dorsal CA2 neurons. However, there is greater molecular heterogeneity in the anatomic vCA2 region compared to its dorsal counterpart, and these neurons project with CA1 with different synaptic strength and feedforward inhibition.

### Avpr1b-expressing cells in vCA2 are not necessary for social memory

Given the well-defined role of dCA2 in social novelty recognition memory, we next examined whether vCA2 played a similar behavioral role (Figure 8). We used a chemogenetic approach, similar to that used previously in examining the role of dCA2^20^, in which we injected a Cre-dependent AAV in either dCA2 or vCA2 of *Avpr1b-Cre* mice to express the hM4Di inhibitory designer receptor exclusively activated by designer drugs (iDREADD) (dCA2 site: Fig. 8b; vCA2 site: Fig. 8e). Cre-littermate controls were injected in dCA2 or vCA2, as appropriate, with the same virus.

**Figure 8.**
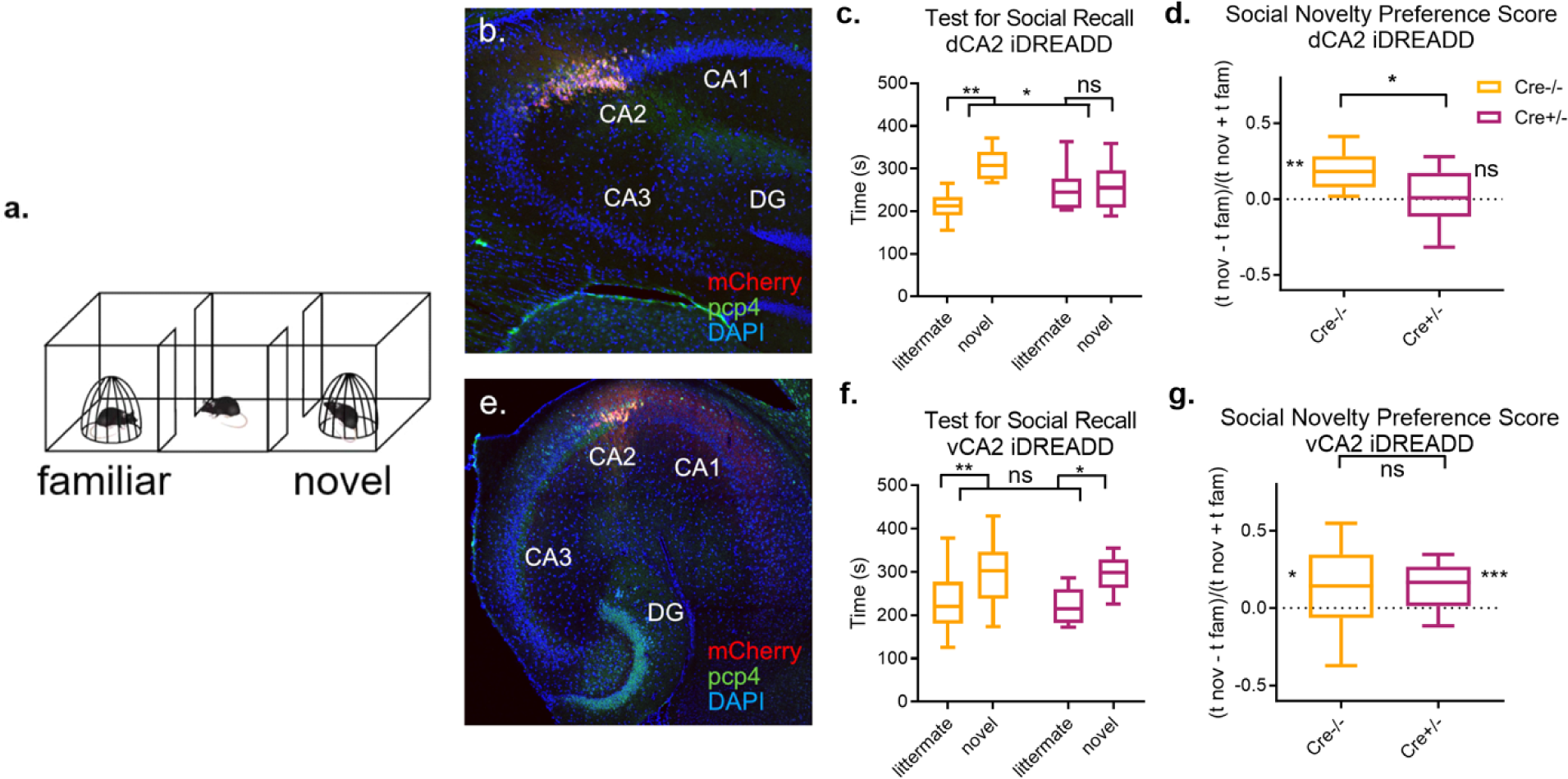
Pharmacogenetic inhibition of Avpr1b-expressing cells in dCA2, but not vCA2, blocks social recall of a littermate. **a)** Test for Social Recall: Following a 10-minute habituation to the three-chamber arena and empty cups, a subject mouse was placed into the arena with a familiar littermate and novel conspecific confined under pencil cups in opposite chambers for 10-minutes. **b)** Site of injection of Cre-dependent mCherry-iDREADD in dCA2 with co-expression of PCP4. **c)** Avpr1b-Cre-/- (control) mice injected with iDREADD in dCA2 (n=10) and administered CNO 30-minutes prior to the test for social recall prefer to explore a novel mouse compared to a familiar littermate. Avpr1b-Cre+/- mice injected with Cre-dependent iDREADD in dCA2 (n=9) and administered CNO 30-minutes prior to the test for social recall spend equal times exploring both individual. Two-way ANOVA Genotype x Interaction Partner F(1,17)=5.617, p=0.030; Šídák’s multiple comparisons test Cre-/- littermate vs. novel p=0.0032, Cre+/- littermate vs. novel p=0.95. **d)** The novelty preference score (the difference in time spent exploring the novel compared to littermate mice, normalized to the total interaction time) is significantly different from zero for Cre-/- mice but not for Cre+/- mice. One-sample t-test against zero: Cre-/- t=4.731, df=9, p=0.0011; Cre+/- t=0.2397, df=8, p=0.82. Unpaired t-test between Cre-/- and Cre+/- t=2.435, df=17, p=0.026. **e)** Site of injection in vCA2 with PCP4 labeling. **f)** Both Cre-/- (n=24) and Cre+/- mice (n=18) interact longer with a novel compared to a familiar littermate. Two-way ANOVA Interaction Partner F(1,40)=16.71, p=0.0002, Genotype x Interaction Partner F(1,40)=0.005, p=0.94; Šídák’s multiple comparisons test littermate vs. novel Cre-/- p=0.0078, Cre+/- p=0.018. g) Both Cre-/- and Cre+/- mice show a significant preference for the novel over the familiar individual. One-sample t-test against zero Cre-/- t=2.648, df=23, p=0.014; Cre+/- t=3.988, df=17, p=0.0010. Unpaired t-test between Cre-/- and Cre+/- t=0.1545, df=40, p=0.88. *p<0.05,**p<0.01, *p<0.05, ns = non-significant.

We first compared the effect of silencing dCA2 and vCA2 in a social novelty recognition task in which a subject mouse is presented with a novel animal and familiar littermate. Three weeks following iDREADD AAV injection, mice were tested for social memory following systemic injection of the iDREADD agonist CNO 30 min prior to testing (Fig. 8). Subjects were first habituated to a three-chamber arena containing empty wire cup cages in the end chambers of the arena. We then placed a familiar littermate and novel conspecific under the wire cups and allowed the subjects to explore the arena for 10 min. Social novelty recognition memory was assessed by the normal preference of a subject to explore the novel individual compared to the familiar littermate. We quantified both the absolute interaction times as well as the behavioral discrimination score, defined as the difference in time spent exploring the novel and littermate mice divided by the total interaction time with both individuals.

As noted previously based on results in the *Amigo2-Cre* mouse line (Hitti and Siegelbaum, 2014), silencing dCA2 fully suppressed the normal preference of the subject to explore the novel compared to familiar mouse. Thus, whereas *Cre-* control mice spent a significantly greater time investigating the novel individual compared to the familiar littermate (Fig. 8c), with a discrimination index significantly greater than zero (Fig. 8d), the *Avpr1b-Cre* group mice failed to distinguish the novel mouse from the familiar littermate. With dCA2 silenced, the subject mice showed no significant difference in interaction time between the novel and familiar individuals, with a discrimination index not significantly different from zero. Thus, in agreement with previous reports in which CA2 was silenced in Amigo2-Cre mice^16,17,20–22,42–46^, we confirmed that dCA2 is crucial for social novelty recognition memory in a distinct mouse line.

In contrast to the effect of silencing dCA2, administration of CNO to *Avpr1b-Cre* mice expressing iDREADD in vCA2 failed to suppress social recognition memory. Both *Avpr1b-Cre* mice and *Cre-* controls spent a significantly greater time interacting with the novel compared to the familiar animal, with a discrimination index significantly greater than zero. There was no significant difference in either interaction times or discrimination index between the two groups of mice (Fig. 8f,g). Thus, in contrast to the importance of dCA2 for social memory, Avpr1b-expressing cells in vCA2 are not necessary for the discrimination of a novel individual from a familiar littermate.

As the ability of an animal to distinguish a novel from familiar littermate primarily relies on recall of a highly consolidated memory, we explored an additional social novelty recognition memory task where subjects must encode and consolidate a new memory of another individual (Suppl. Fig. 9a). In this task, a subject mouse first explored two novel individuals under pencil cup cages in an open arena for 5 min. Following this learning trial, the subject was removed from the arena and returned to its home cage. After 30 min, the subject was returned to the arena for a 5 min recall trial in which the subject again explored two mice in wire cup cages, with one mouse from the learning trial and the other a previously unencountered novel individual. Social recognition memory was assessed by the preferential exploration of the novel individual compared to the now familiarized individual previously encountered in the learning trial.

Previous studies have shown that silencing dCA2 with chemogenetics or optogenetics during the learning trial prevents the discrimination of the novel and familiarized mouse in the recall trial^20,21^. In contrast, we found that systemic injection of CNO 30 min prior to the social memory test in Avpr1b-Cre mice expressing hM4Di in vCA2 failed to inhibit social memory. Thus, both Avpr1b-Cre mice and Cre-control littermates (both injected with CNO and Cre-dependent AAV) spent significantly more time interacting with the novel individual compared to the more familiar one (Suppl. Fig. 9b), with discrimination indexes significantly greater than zero (Suppl. Fig. 9c). Again, no difference was observed between groups for either measure (Two-way repeated measures ANOVA Genotype x Interaction Partner F(1,38)=0.6985, p=0.70; unpaired t-test t=0.1270, df=38, p=0.90. These results confirm the lack of social memory deficit we observed in the littermate recognition test, and further confirm that Avpr1b-expressing cells in vCA2 are not necessary for social memory encoding or recall.

Because the *Avpr1b-Cre* line leads to Cre expression in a smaller fraction of vCA2 neurons (38.0 ± 5.4%) compared to dCA2 neurons (72.4 ± 8.8%), it is possible that the lack of effect of silencing vCA2 on social memory is due to the lower proportion of total cells in vCA2 that are inhibited with this line. As we found that PCP4 is expressed in a wider fraction of vCA2 neurons compared to Avpr1b-Cre-mediated expression (60% compared to 38%), we next determined the efficacy of chemogenetic silencing of vCA2 on social memory using a PCP4-Cre mouse line.

As this line has not been previously used to target CA2, we first examined the efficacy and specificity of its ability to target vCA2 (Suppl. Fig. 10a-b). We found that injection of Cre-dependent AAV in vCA2 led to widespread and relatively selective expression in vCA2 (Fig. 5a). However, as PCP4 is also expressed in dentate gyrus, extra care was taken with the targeting of viral injection to limit expression to CA2. Quantitatively, 53.86% ± 2.3% (n=16) DAPI-expressing cells were infected with Cre-dependent virus in vCA2, which is close to the proportion of cells that express PCP4 in vCA2.

We performed the same social memory recall test with a novel animal and familiar littermate we used for the *Avpr1b-Cre* line, and again found that silencing the broader population of PCP4-expressing vCA2 neurons did not alter social recall (Suppl. Fig. 10c-d). Thus both *PCP4-Cre* and *Cre-* littermates showed normal discrimination of the novel individual from the familiar littermate, with no significant difference between groups (Two-way repeated measures ANOVA, Genotype x Interaction Partner F(1,17)=0.7533, p=0.75; 32.20; Šídák’s multiple comparisons test for exploration of littermate versus novel mouse: Cre-control, p=0.0024; PCP4-Crre, p=0.0014; Suppl. Fig 10c-d). Altogether these results suggest that vCA2 neurons are less critical for social memory compared to dCA2.

### PCP4-expressing cells in vCA2 modulate social aggression

In addition to its role in social memory, dCA2 promotes social aggression^34,40^ As ventral hippocampus has been implicated in behaviors tied to emotions, we hypothesized that vCA2 may play a more important role in regulating aggression than in social memory discrimination. We thus used the resident-intruder test to determine the effect of silencing vCA2 on aggression in PCP4-Cre residents expressing iDREADD compared to Cre-controls (Figure 9). Three weeks following viral injection, we introduced a novel BALBc intruder into the cages of singly housed residents for 5 min and repeated this test once a day over three successive days using the same BALBc intruder. On days 1-3 we injected both cohorts of resident mice with saline 30 min prior to testing aggression. On day 4 we examined the effect of silencing CA2 on aggression by injecting CNO systemically in both groups of mice using a novel BALBc intruder for a 10-min resident-intruder test. PCP4-Cre and Cre-saline-injected residents mounted similar levels of aggression to the BALBc intruder on days 1-3, measured either by the number of biting attacks or total attack duration, indicating that there was no effect of genotype on aggression. In contrast, injection of CNO led to a large decrease in aggression towards the BALBc intruder in the iDREADD-expressing *PCP4-Cre* mice relative to the *Cre-* control group (Figure 9). CA2 silencing caused a significant decrease in both the total duration of aggression (median = 12.3 s for *Cre-* and 0.15 s for *PCP4-Cre* groups, Mann Whitney test, p=0.0042) and the number of bouts of aggression (median = 13 for *Cre-* and 0.5 for *PCP4-Cre* groups, Mann Whitney test, p=0.018). Thus, although vCA2 does not appear to be required for normal social memory behavior, it is necessary to promote social aggression, similar to the role of dCA2.

**Figure 9.**
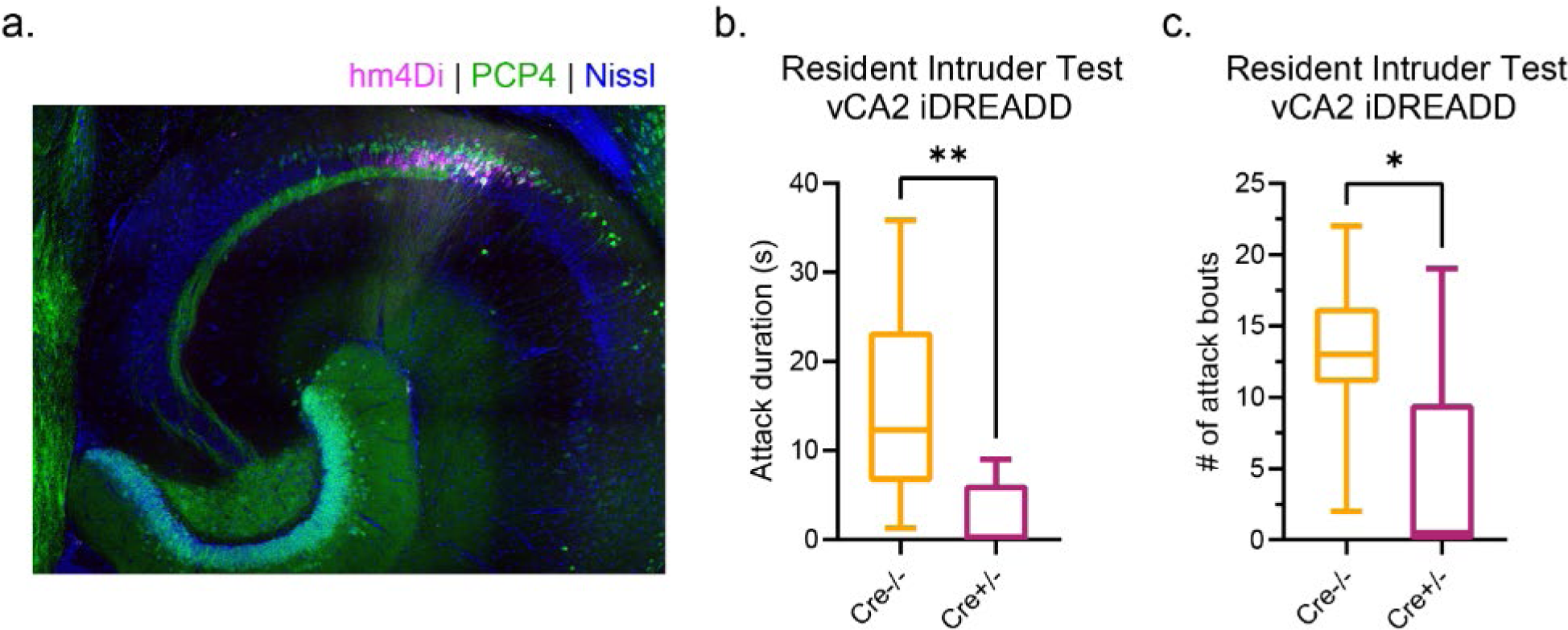
Inhibition of vCA2 PCP4-expressing neurons decreases social aggression. **a)** Site of injection of Cre-dependent mCherry-iDREADD in vCA2 with co-expression of PCP4 in a PCP4-Cre+/- subject animal. Scale bar, 200 µm. **b)** Three weeks following viral injection, subjects were injected with CNO and exposed to a novel mouse in a resident-intruder test. Time spent engaged in social attack was quantified, revealing a significant decrease in Cre+/- subjects (n=8) compared to controls (n=8) (median tCre-/- = 12.3 s, median tCre+/- = 0.15s, Mann Whitney test p=0.0042). **c)** Cre-/- subjects have significantly more attack bouts compared to Cre+/- subjects injected with iDREADD and CNO (median Cre-/- = 13 attacks, median Cre+/- = 0.5 attacks, Mann Whitney test p=0.018). **p<0.01, *p<0.05

## Discussion

Here we provide the first comprehensive analysis of the molecular, morphological, and electrophysiological properties of ventral CA2 pyramidal neurons and examine their behavioral role. Our results show the existence of populations of pyramidal neurons in the anatomically defined CA2 region in ventral hippocampus that share certain characteristic molecular markers, cellular morphology, and electrophysiological and synaptic properties with the well-defined population of CA2 neurons in dorsal hippocampus. However, unlike the relative uniform pattern of molecular marker expression among dorsal CA2 pyramidal neurons, we find distinct subpopulations of pyramidal neurons based on molecular marker expression in ventral CA2. Moreover, we show that ventral CA2 pyramidal neurons show both similar and distinct behavioral roles compared to dorsal CA2.

Like dorsal CA2 pyramidal neurons, ventral CA2 neurons have a characteristically low input resistance, high capacitance and low sag ratio, properties that distinguish them from ventral CA1 pyramidal neurons. Moreover, in contrast to CA3 pyramidal neurons, ventral CA2 neurons, by and large, lack thorny excrescences. As in dorsal CA2, ventral CA2 neurons receive a very strong excitatory drive from their direct entorhinal cortex inputs that target the distal apical dendrites in stratum lacunosum moleculare. Also similar to dorsal CA2, the major intrahippocampal output from ventral CA2 targets CA1 and the major extrahippocampal output from ventral CA2 targets the lateral septum.

However, whereas the vast majority of dorsal CA2 pyramidal neurons co-express a number of characteristic molecular markers, including the Avpr1b receptor, RGS14, and PCP-4, there are at least two subpopulations of molecularly distinct pyramidal within vCA2. One subpopulation, representing about 40% of the neurons in the CA2 region, is largely confined to the superficial layer of vCA2 (near the border with stratum radiatum) and co-expresses Avpr1b, PCP4 and RGS14. A second subpopulation, representing about 30% of vCA2 neurons, does not express Avpr1b or RGS14 but does express PCP4. A large fraction of the remaining 30% of neurons that express neither Avpr1b nor PCP4 likely correspond to inhibitory neurons, although the exact fraction remains to be determined. It is also important to note that the markers we examined here are just a small fraction of the proteins that characterize dorsal CA2, and there may be unique markers of ventral CA2 that have not yet been identified. Thus further work is needed to fully characterize the cells in this region, perhaps through the extension of RNA sequencing profiles that have explored expression from other subregions of the hippocampus^3^.

Our results also provide important insights into the behavioral roles of ventral CA2 pyramidal neurons and their similarities and differences with dorsal CA2. Unlike dorsal CA2 pyramidal neurons, neither the Avpr1b-expressing subpopulation nor the larger PCP4-expressing subpopulation of ventral CA2 pyramidal neurons are necessary for social novelty recognition memory. In contrast, we find that the PCP4-expressing ventral CA2 cells are necessary for the promotion of social aggression, similar to prior behavioral findings on the importance of dorsal CA2 pyramidal neurons^34,40,47^. These results are consistent with the view that dorsal hippocampus is more important for cognitive function whereas ventral hippocampus is more important for regulating emotional behaviors, such as anxiety.

The differences and similarities in the behavioral roles of ventral compared to dorsal CA2 may reflect the differences and similarities in the synaptic targets of these two regions. Our laboratory previously found that dCA2 regulates social novelty recognition memory through its projections to vCA1a^20^, which selectively target the stratum radiatum region of distal vCA1 (vCA1a), bordering the subiculum. In contrast, we here found that vCA2 targets the stratum oriens of a complementary region of regions of vCA1 more proximal to vCA2 (vCA1b and vCA1c). Thus, the differential roles of dCA2 and vCA2 in social novelty recognition memory may be a result of different synaptic targeting within vCA1.

A second possible reason as to why vCA2 pyramidal neurons are not required for social novelty recognition memory, which is suggested by our results, is that the vCA2 projections to vCA1a elicit a very large feedforward inhibition, which is greater than the inhibitory response evoked by activation of the projections from dCA2 to dCA1. Thus, activation of dCA2 produced a significantly smaller IPSC in the deep-layer neurons in dCA1 than the IPSC evoked in vCA1 in response to activation of its vCA2 inputs. Thus, we propose that vCA2 serves to dampen, rather than excite, vCA1a neuron activity, in opposition to the excitatory action of dCA2, consistent with our finding that silencing vCA2 does not inhibit social recognition memory. It remains unclear what role, if any, vCA2 plays in other forms of hippocampal-dependent memory or emotional behavior.

In contrast to their differential roles in regulation social memory, vCA2 and dCA2 exert a similar action to promote social aggression. Of particular interest, both regions appear necessary to mount an aggressive response as inhibition of either region alone is sufficient to almost suppress aggressive behavior fully. While we have not identified which vCA2 projections serve to modulate social aggression, we hypothesize that the vCA2 output to lateral septum is most relevant, based on prior studies showing that dCA2 pyramidal neurons promote aggression through their projections to dorsal lateral septum^40^. Such projections act by exciting GABAergic neurons in dorsal lateral septum that project to and inhibit GABAergic neurons in ventral lateral septum that normally provide tonic inhibitory input to neurons that trigger socially aggressive behavior in the ventrolateral subdivision of the ventromedial nucleus of the hypothalamus (VMHvl)^48^. As a result, dCA2 activation disinhibits VMHvl, thereby promoting aggression. Although vCA2 targets a region of lateral septum that is more ventral than that targeted by dCA2, both projection zones fall in the confines of dorsal lateral septum. We thus hypothesize that, similar to dCA2, vCA2 projections to lateral septum lead to disinhibition of VMHvl. To explain how silencing of either dCA2 or vCA2 can markedly suppress aggression, we hypothesize that the two sets of dorsal lateral septum neurons activated by dCA2 and vCA2 may converge on a common population of ventral lateral septum neurons, acting synergistically to cause sufficient inhibition of ventral lateral septum output needed to disinhibit VMHvl.

Previous studies have demonstrated the importance of Avpr1b, which is highly enriched in CA2 pyramidal neurons compared to almost all other regions of the brain, in both promoting social aggression (Wersinger, 2002; Pagani, 2015; Leroy, 2018) and social memory behavior (Smith, 2016). Pagani et al. found that social aggression behavior could be partially rescued in a general Avpr1b knockout mice by lentiviral-mediated expression of Avpr1b targeted mainly to dorsal CA2. Of interest, although viral expression of Avpr1b in dCA2 restored to normal levels the percentage of mice that attacked an intruder, attack latency was only partially rescued. Moreover, attack duration and number of attacks were two-fold lower in the rescued group compared to wild-type mice, although the difference was not quite statistically significant (p=0.07). This incomplete rescue could be explained if Avpr1b expression in vCA2 was necessary to achieve normal levels of aggression.

At present it is uncertain as to whether the subpopulation of PCP4-expressing neurons in vCA2 that do not express Avpr1b contribute to aggression. However, it is of interest that the selective rescue of expression of Avpr1b in dCA2 (but not vCA2) in the Avpr1b knockout mice (Pagani 2015) restores aggression to a level greater than that seen in our experiments when we selectively silenced PCP4-expressing vCA2 neurons (which caused a near complete loss of aggression; see Fig.9). These combined results indicate that vCA2 must normally contribute to aggression in a manner independent of Avpr1b. This action could either reflect a contribution of Avpr1b-expressing vCA2 neurons that does not require Avpr1b activation and/or it could represent a contribution of the PCP4-expressing vCA2 neurons that do not express Avpr1b.

Altogether, our results show the existence of a broadly defined category of CA2 pyramidal neurons that share certain molecular, anatomical and electrophysiological properties along the entire longitudinal axis of the hippocampus. However, dorsal and ventral CA2 regions show important differences in molecular characteristics, synaptic output connectivity, and roles in regulating social memory and behavior. Whereas dCA2 contributes both to social novelty recognition memory and social aggression, vCA2 is not required for social memory and is more selectively involved in promoting social aggression. Future work is needed to assess whether the lack of involvement of vCA2 in social novelty recognition memory is because vCA2 neurons fail to encode the type of information about social novelty/familiarity that is present in dCA2^44,46^. Alternatively, novelty and familiarity may be encoded in vCA2 similarly to dCA2 but the output from the subpopulation of vCA1 neurons that receive input from vCA2 do not target downstream brain regions that participate in social novelty recognition memory behaviors. If, as hypothesized, vCA2 modulates social aggression through its output to lateral septum, the purpose of its prominent projections within the hippocampus to vCA1b and vCA1c remains mysterious.

## Supplemental Figures

**Supplementary Figure 1.**
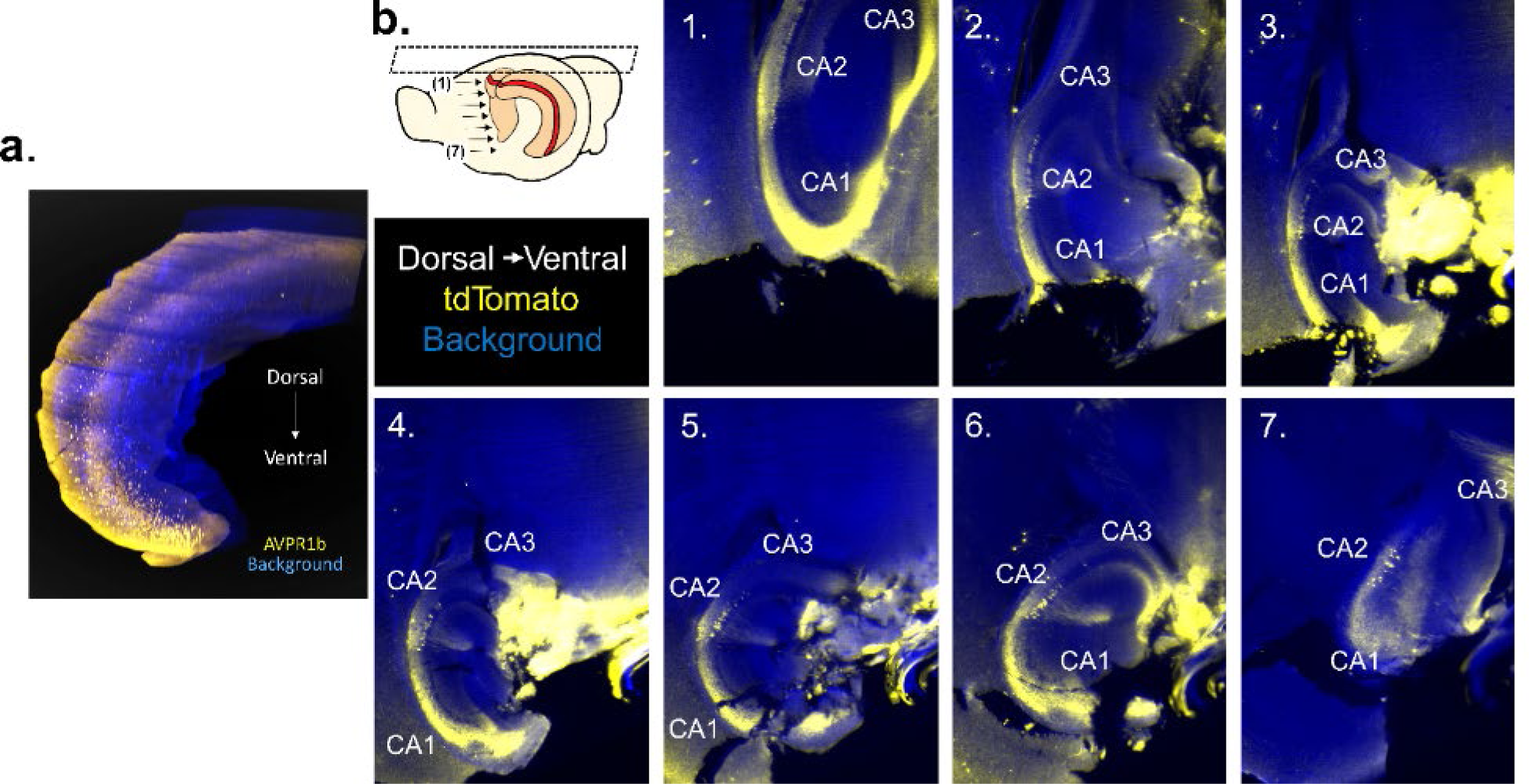
Light sheet microscopy of iDISCO-cleared Avpr1b-Cre x Ai14 brains (n=3) confirms expression of Avpr1b along the entire dorsoventral axis. **a)** Composite image of the entire hippocampus. **b)** Individual sheets from most dorsal (1) to most ventral (7).

**Supplementary Figure 2.**
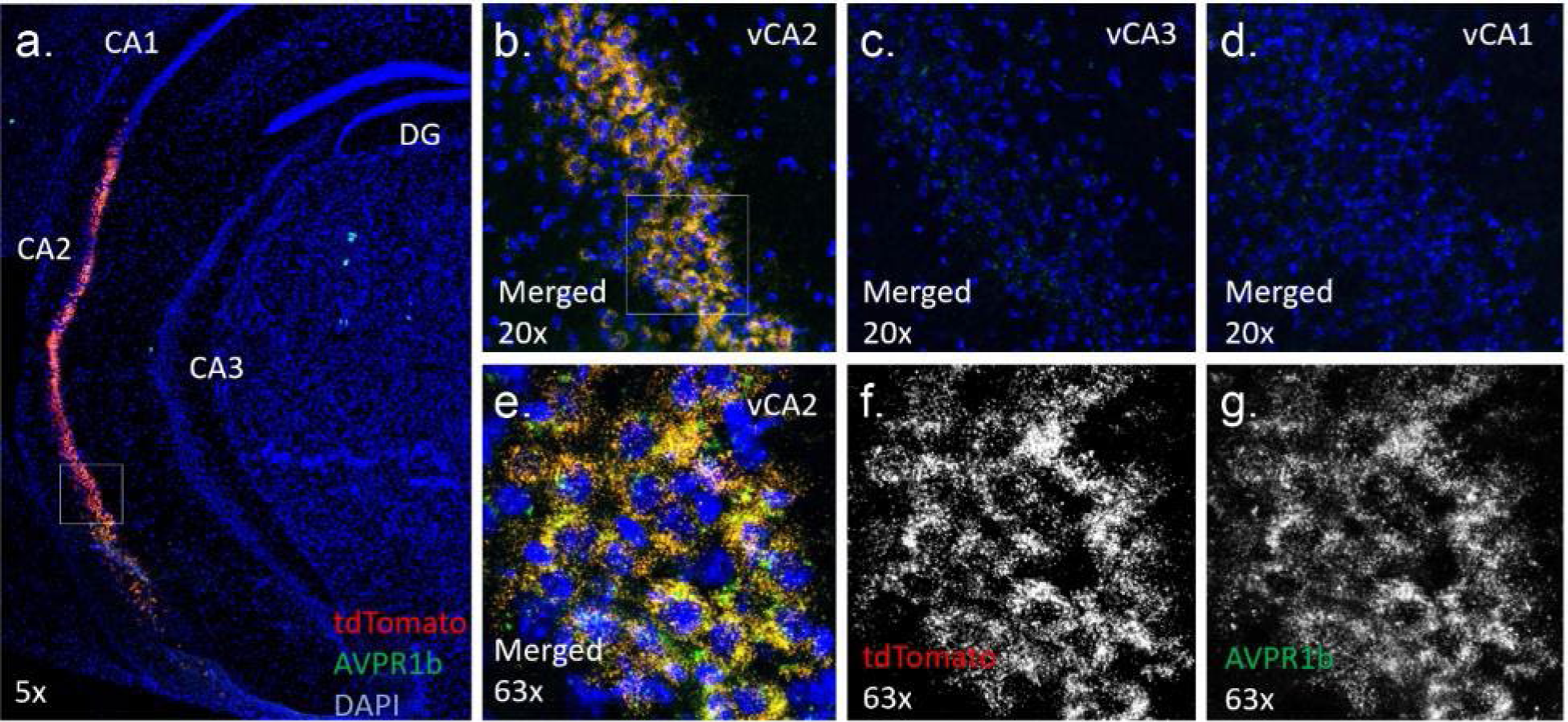
In-situ hybridization of Avpr1b and tdTomato shows co-expression along the entire dorsoventral axis in Avpr1b-Cre x Ai14 mouse brains (n=2). **a)** 5x coronal slice showing Avpr1b and tdTomato co-expression. **b)** 20x image of vCA2 in coronal slice. No Avpr1b hybridization is observed in **c)** ventral CA3 or **d)** ventral CA1. **e)** 63x image of vCA2 with expression of tdTomato (red) and Avpr1b (green). **f)** Red channel (tdTomato). **g)** Green channel (Avpr1b).

**Supplementary Figure 3.**
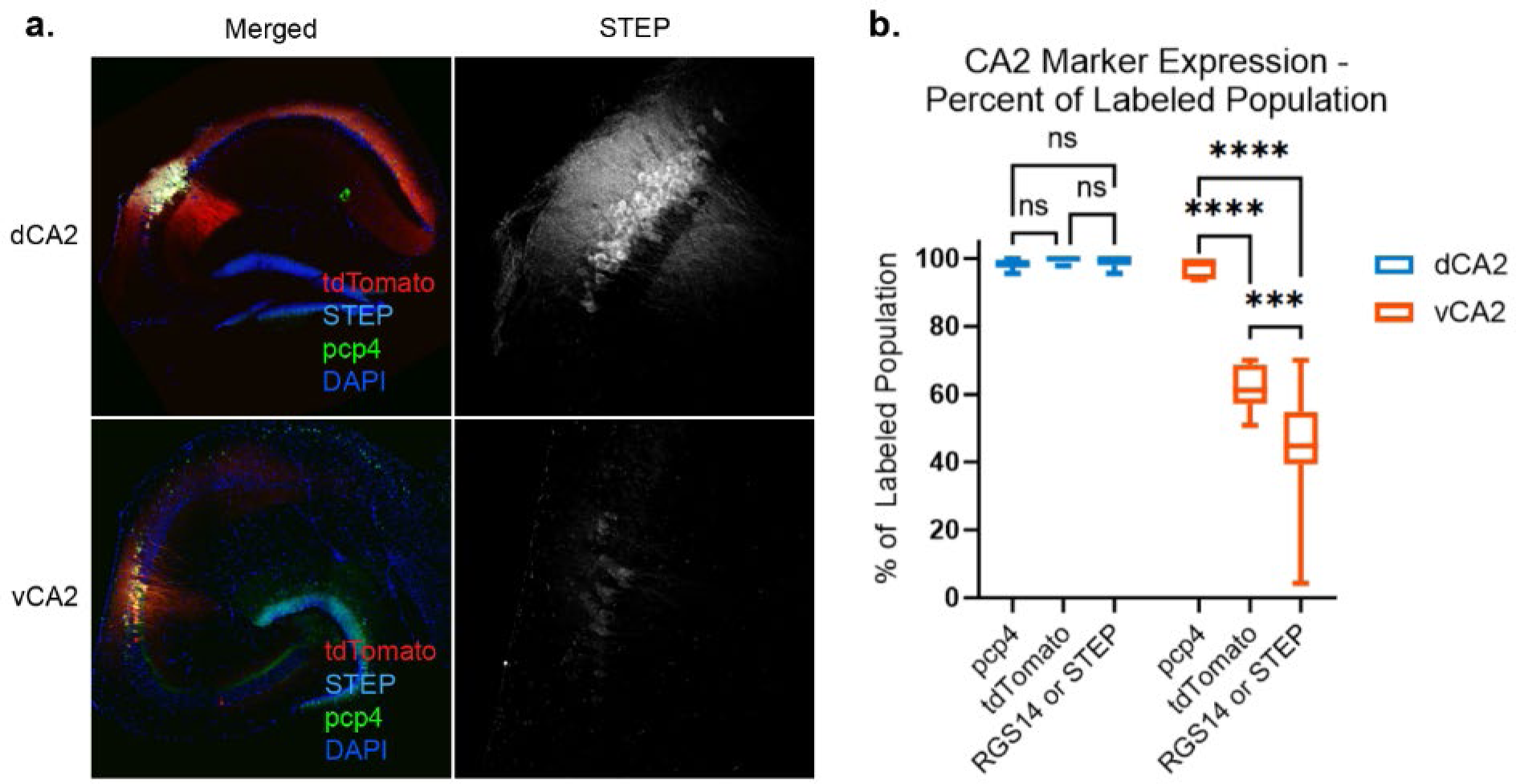
Quantification of dorsal and ventral neurons expressing PCP4, tdTomato (Avpr1b), or RGS14/STEP. **a)** Slices from Avpr1b-Cre x Ai14 mice show reduced STEP expression in vCA2 compared to dCA2. **b)** There is greater heterogeneity of expression in vCA2 compared to dCA2 of cells expressing PCP4, tdTomato, and RGS14/STEP (Two-way ANOVA Dorsoventral Region x Protein F(2,39) = 39.21, p<0.0001. Tukey’s multiple comparisons test: dCA2 PCP4 vs. dCA2 tdTomato p=0.96, dCA2 PCP4 vs. dCA2 RGS14/STEP p=0.98, dCA2 tdTomato vs. dCA2 RGS14/STEP p=0.99, vCA2 PCP4 vs. vCA2 tdTomato p<0.0001, vCA2 PCP4 vs. vCA2 RGS14/STEP p<0.0001, vCA2 tdTomato vs. vCA2 RGS14/STEP p=0.0007. ns = non-significant, *** p<0.001, **** p<0.0001.

**Supplementary Figure 4.**
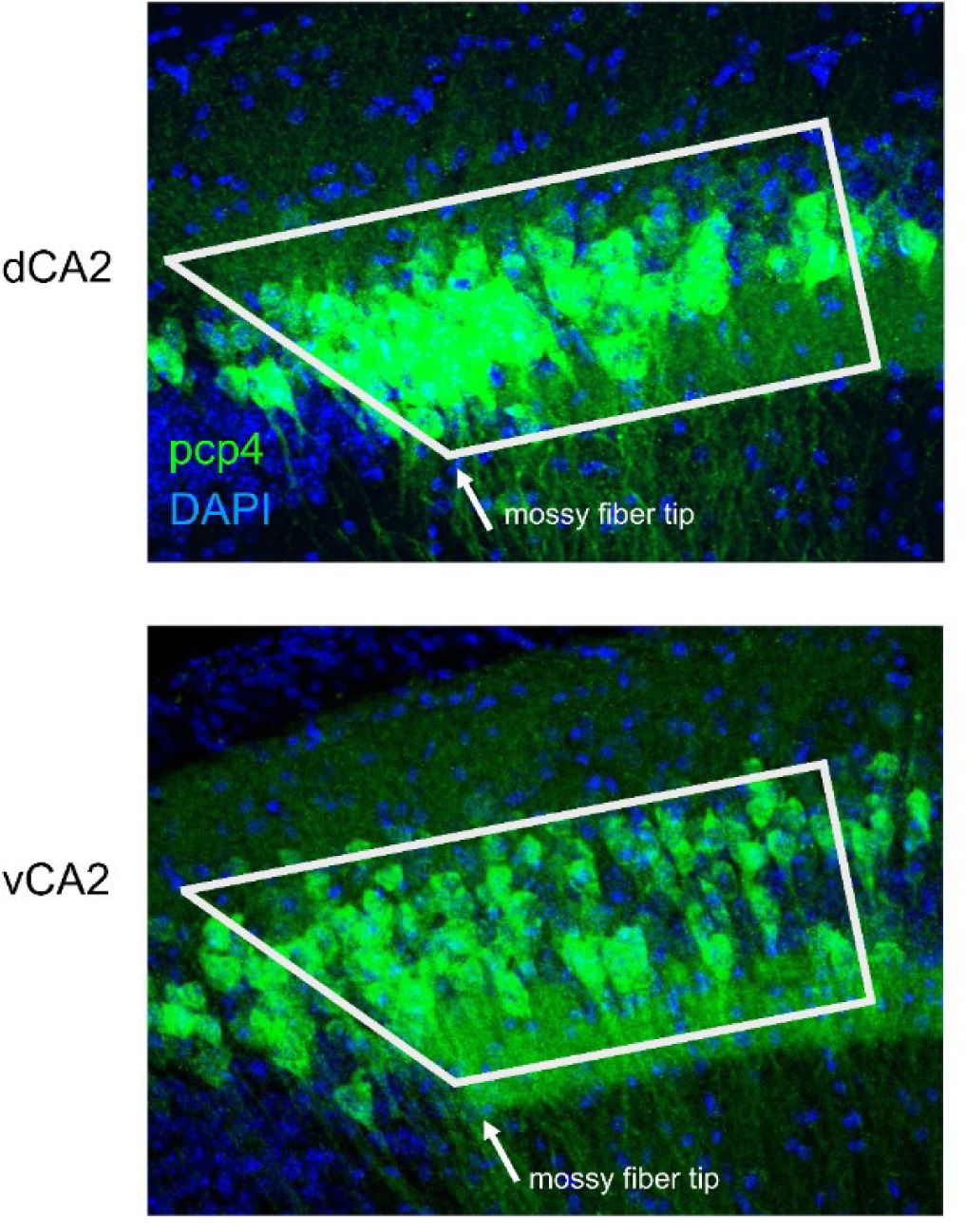
dCA2 and vCA2 are located at the tip of the PCP4-labeled mossy fiber. Transverse hippocampal slices show PCP4- and DAPI-labeled dCA2 (top) and vCA2 (bottom). Images intentionally over-exposed to highlight PCP4-labeled mossy fiber (tip of pcp4-labeled mossy fiber denoted by white arrow). White box shows standardized central region of CA2 captured by cell counting.

**Supplementary Figure 5.**
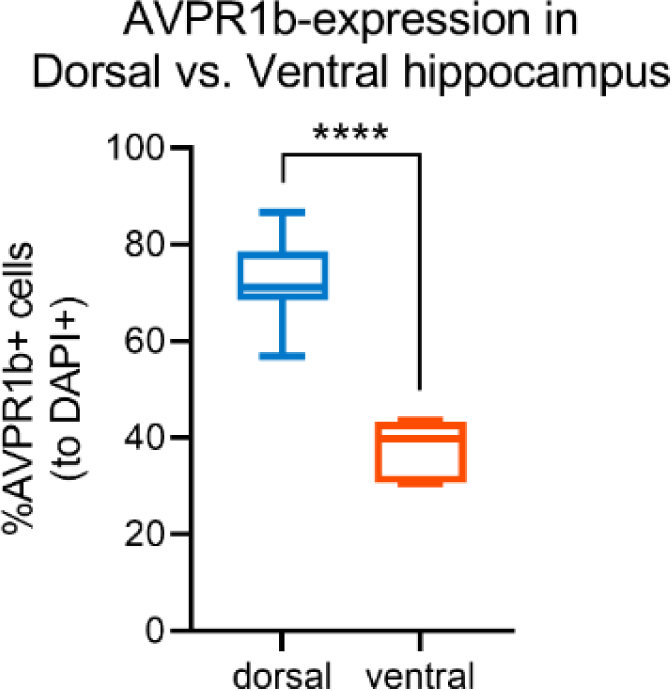
Quantification of tdTomato (AVPR1b) expression in dorsal and ventral transverse hippocampal slices. A significant number of cells express tdTomato (AVPR1b) in both dorsal (n=8) and ventral (n=7) slices from AVPR1b-Cre x Ai14 mice (dorsal: 72.4 ± 8.8%, one-sample t-test against zero t=23.25, df=7, p<0.0001; ventral: 38.0 ± 5.4%, one-sample t-test against zero t=18.62, df=6, p<0.0001). Significantly fewer cells express tdTomato (AVPR1b) in the ventral hippocampus compared to the dorsal hippocampus (two-sample unpaired t-test: t=8.953, df=13, p<0.0001).

**Supplementary Figure 6.**
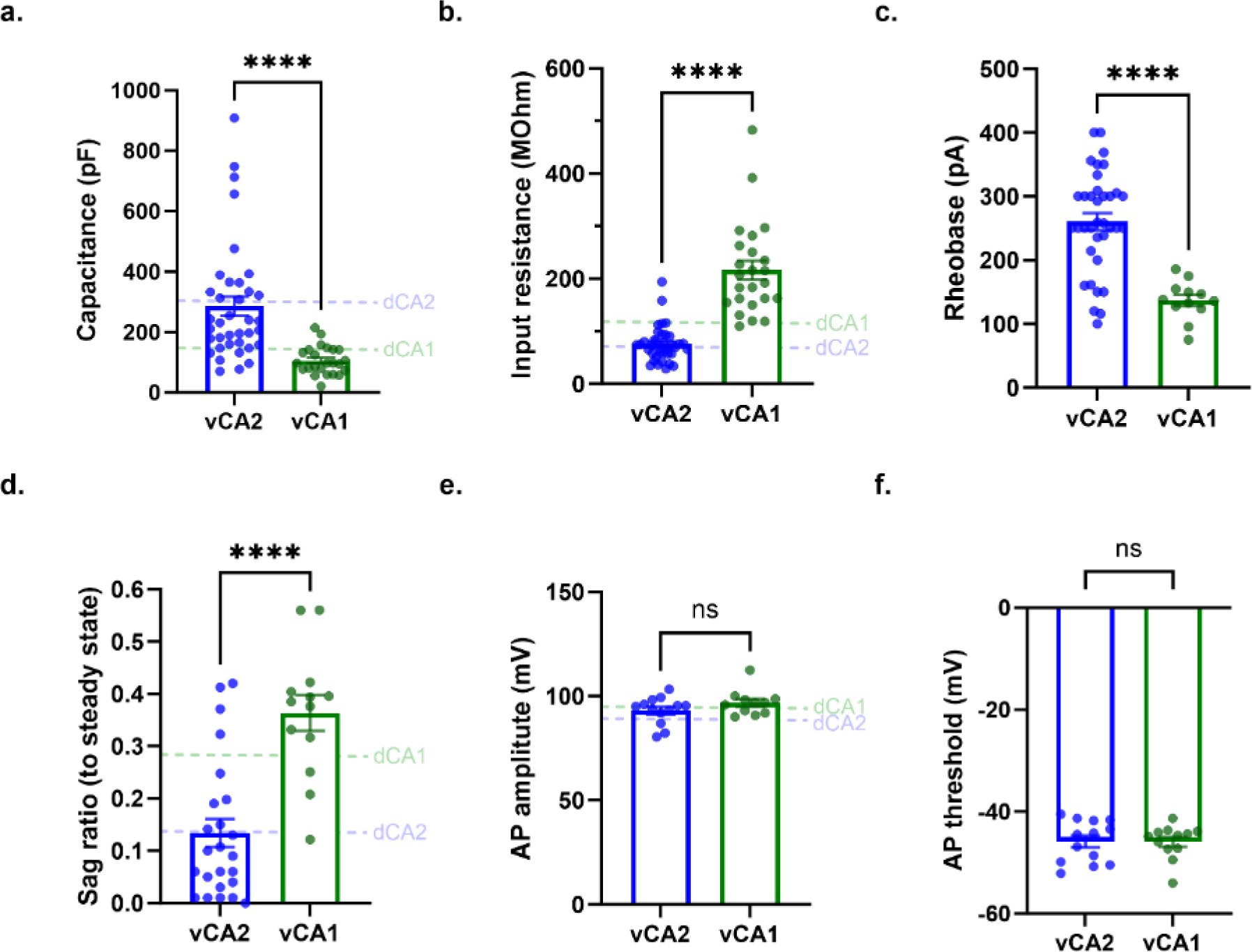
Intrinsic properties of vCA2 neurons compared to vCA1 neurons. **a)** vCA2 has significantly higher capacitance compared to vCA1 (Unpaired t test with Welch’s correction: t=5.411, df=42.09, p<0.0001, vCA2: n_cell_=37, n_mouse_=10, vCA1: n_cell_=24, n_mouse_=7). Membrane capacitance was obtained at −70mV from a V-shape voltage clamp ramp protocol with ΔV=±10mV. **b)** vCA2 has significantly lower input resistance compared to vCA1 (Unpaired t test with Welch’s correction: t=7.489, df=27.32, p<0.0001, vCA2: n_cell_=37, n_mouse_=9, vCA1: n_cell_=24, n_mouse_=7). Input resistance was calculated from a voltage step from −70mV to −80mV in voltage clamp configuration. **c)** vCA2 has significantly higher rheobase compared to vCA1 (Unpaired t test with Welch’s correction: t=7.856, df=44.64, p<0.0001, vCA2: n_cell_=36, n_mouse_=, vCA1: n_cell_=12, n_mouse_=4). **d)** vCA2 has significantly lower sag ratio compared to vCA1 (Unpaired t test: t=5.165, df=35, p<0.0001, vCA1: n_cell_=13, n_mouse_=6). **e)** There is no significant difference in AP amplitude in vCA2 or vCA1 neurons (Unpaired t test: t=1.395, df=22, p=0.1769, vCA2: n_cell_=13, n_mouse_=9, vCA1: n_cell_=11, n_mouse_=4). **f)** There is no significant difference in AP threshold between vCA2 and vCA1 (Unpaired t test: t=0.03140, df=23, p=0.9752, vCA2: n_cell_=13, n_mouse_=9, vCA1: n_cell_=12, n_mouse_=4). Intrinsic properties of dCA2 and dCA1 neurons (dashed lines) are adapted from Chevaleyre and Siegelbaum (2010). ****p<0.0001, ns = non-significant

**Supplementary Figure 7.**
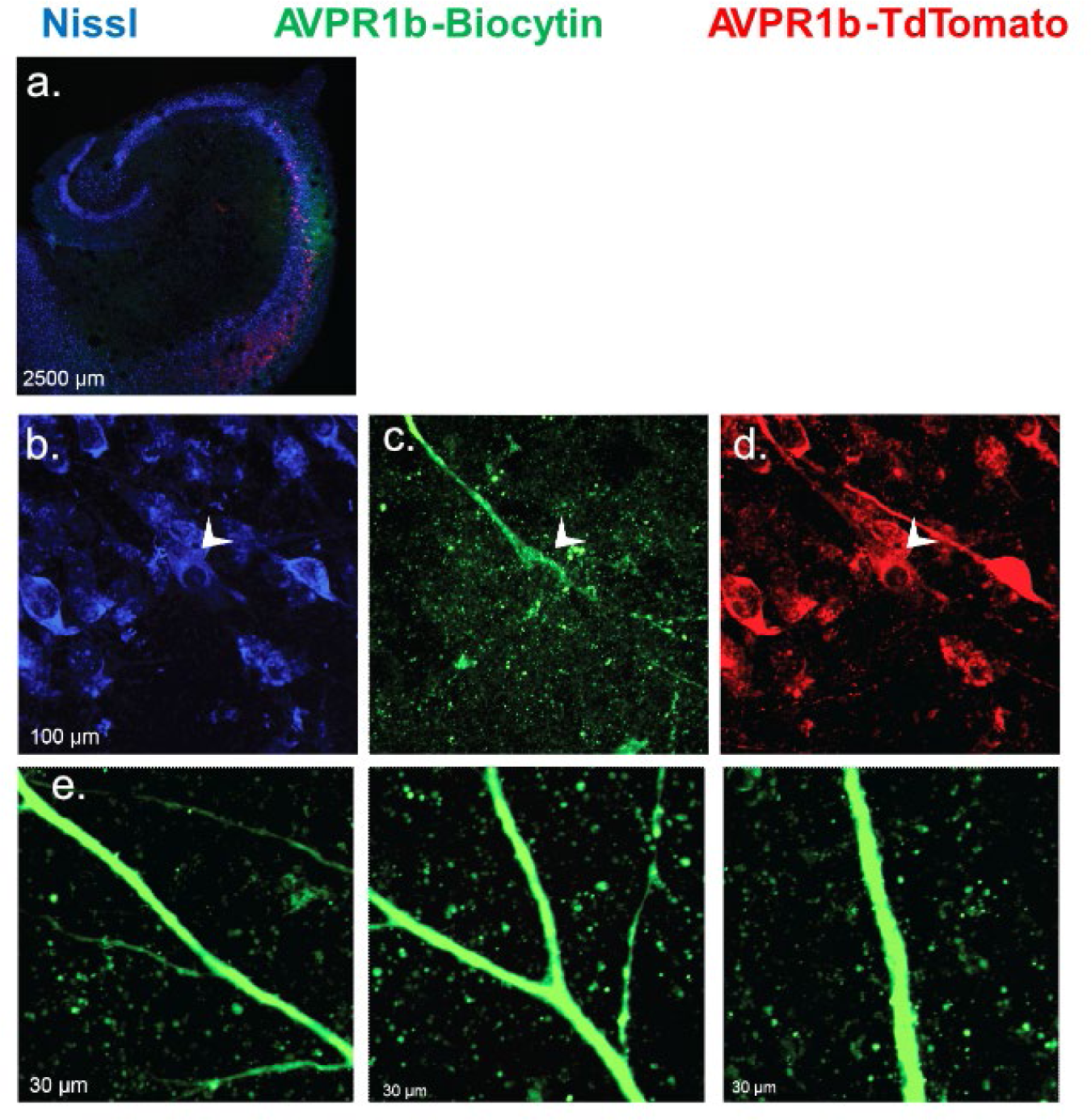
Biocytin filling of cells labeled with tdTomato in Avpr1b-Cre x Ai14 transverse slices reveals these neurons lack thorny excrescences. **a)** Example transverse slice from ventral hippocampus. **b)** Nissl staining. **c)** Avpr1b-expressing cell filled with biocytin. **d)** Avpr1b-expressing cell showing tdTomato expression. **e)** Example zoomed images of Biocytin-filled processes from Avpr1b-expressing neurons showing absence of thorny excrescences. Bottom left corner indicates the number of pixels in image width.

**Supplementary Figure 8.**
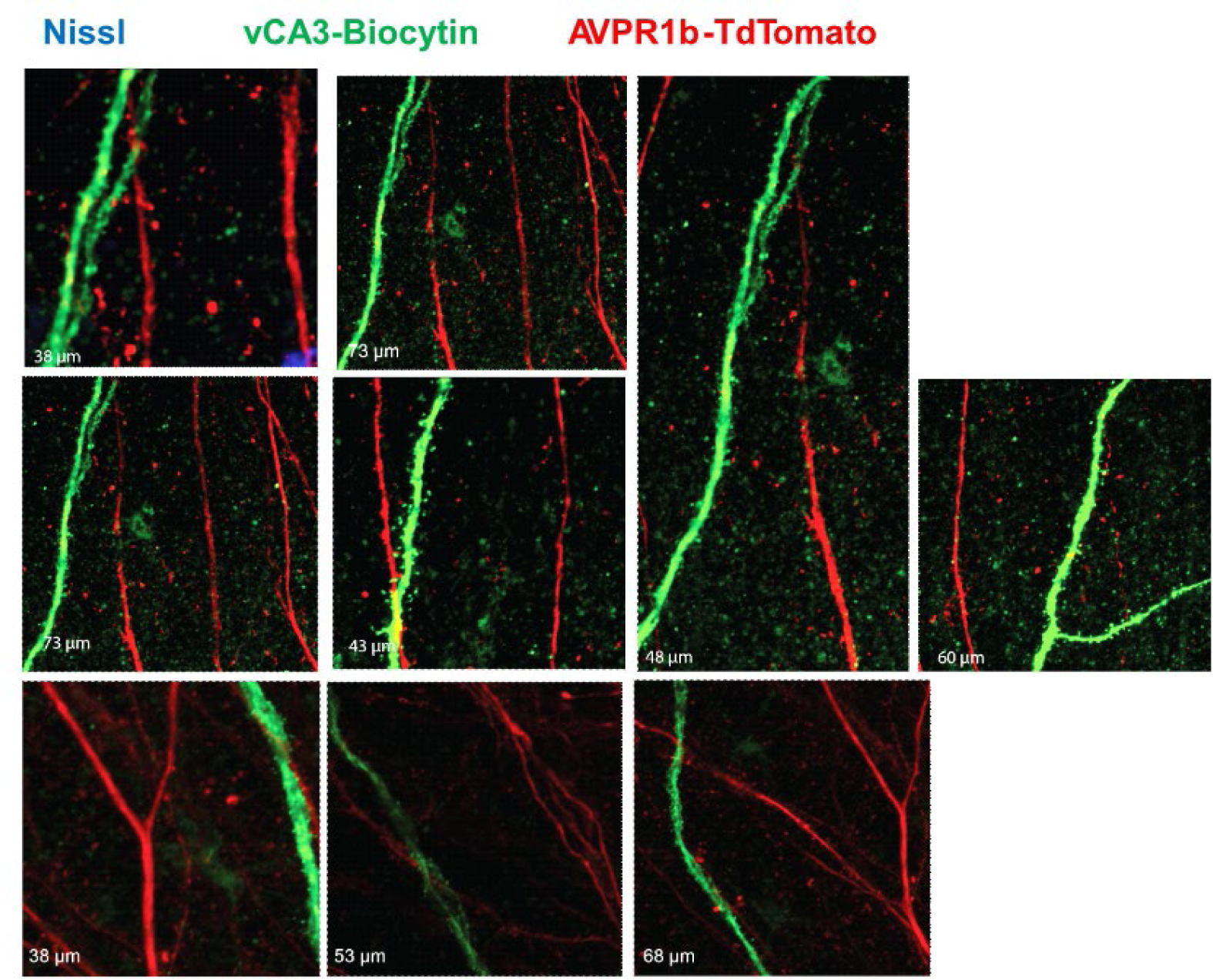
vCA3 cells labeled with biocytin from Avpr1b-Cre x Ai14 transverse slices show thorny excrescences, while cells that express tdTomato (Avpr1b) do not. vCA3 neuron labeled with biocytin does not express tdTomato and shows thorny excrescences from synapses with dentate gyrus. vCA2 neuron labeled with tdTomato does not show thorny excrescences. Bottom left corner indicates the number of pixels in image width.

**Supplementary Figure 9.**
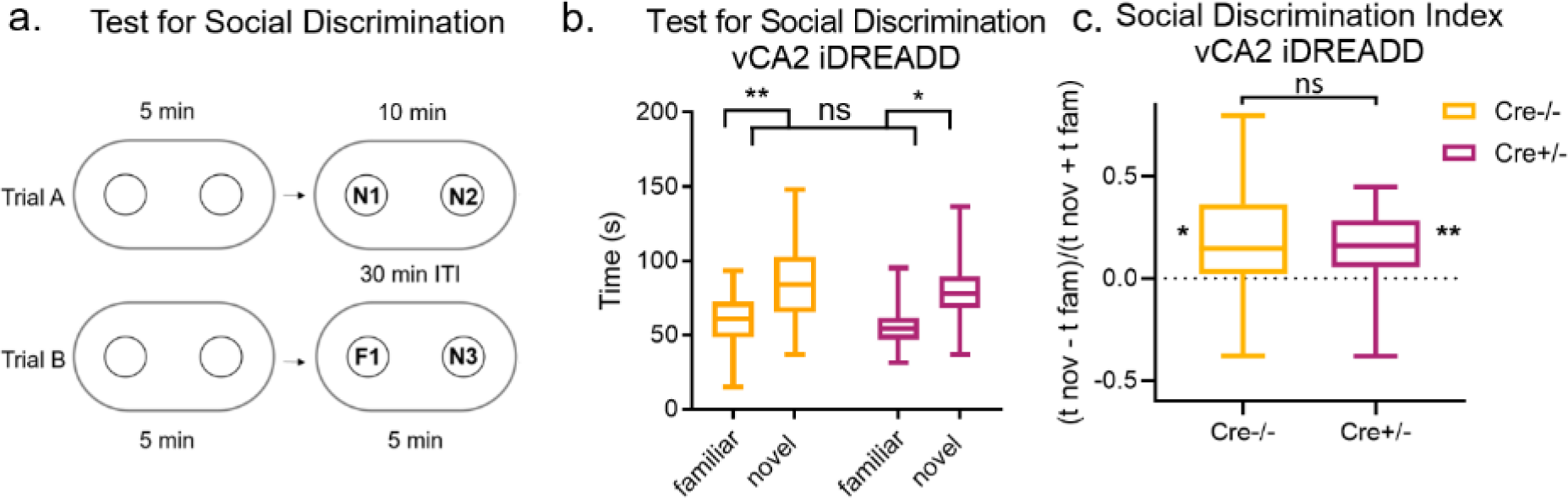
Pharmacogenetic silencing of Avpr1b-expressing vCA2 pyramidal neurons does not impact social recognition. **a)** Avpr1b-Cre-/- and Cre+/- mice were injected with an Cre-dependent mCherry-iDREADD virus in vCA2. On the experiment day, mice were injected intraperitoneally with CNO and following a 30-minute period were run through the Test for Social Discrimination. The test consists of two trials with two investigation periods. In the first period of trial A, mice are habituated to empty cups. In the second trial, mice are exposed to two novel individuals (N1 and N2) under pencil cups in an oval arena, with an individual at either end of the arena, for ten minutes. Following a 30-minute intertrial interval, mice are again habituated to empty cups. Finally, they are exposed to two individuals under pencil cups: one of the now-familiar mice from trial A (F1), and a third novel mouse (N3). **b)** Cre-/- (n=22) and Cre+/- (n=18) mice prefer to investigate the novel individual over a familiar individual (Two-way repeated-measures ANOVA Interaction Partner F(1,38)=17.20, p=0.0002; Šídák’s multiple comparisons test familiar vs. novel Cre-/- p=0.0034, Cre+/- p=0.031). **c)** Cre-/- and Cre+/- mice show a significant preference to interact with the novel over the familiar individual (one-sample t-test against zero Cre-/- t=2.793, df=21, p=0.011; Cre+/- t=3.158, df=17, p=0.0057).

**Supplementary Figure 10.**
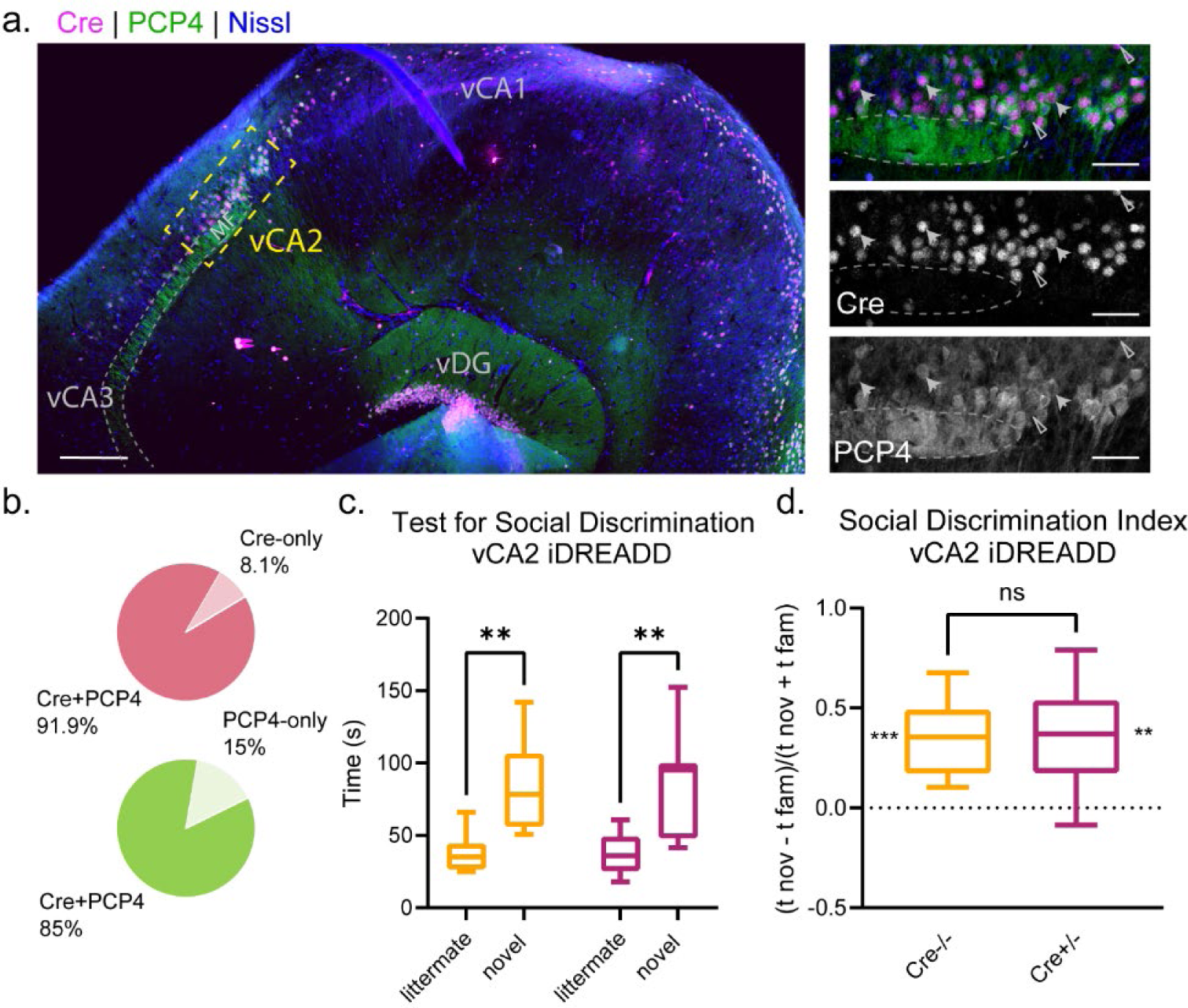
Pharmacogenetic silencing of PCP4-expressing vCA2 pyramidal neurons does not impact social recognition. **a)** A representative transverse ventral hippocampal slice stained for Cre (magenta) in a PCP4-Cre+/- mouse demonstrating co-expression with PCP4 antibody staining (green). Mossy fiber (MF) ends from DG were circled with dashed lines. Left: low magnification view of the ventral hippocampal slice. Scale bars, 200 µm. Right: high magnification view of vCA2. Solid arrows point example co-labeled cells, while open arrows point example cells that are labeled with only Cre of PCP4. Scale bars, 50 µm. **b)** Quantification of the expression of Cre and PCP4 in vCA2 slices (n=2). **c)** PCP4-Cre-/- (n=10) and PCP4-Cre+/- mice (n=8) were both injected with iDREADD virus. Following three weeks, subjects were injected intraperitoneally with CNO and following a 30-minute period were run through the test for social discrimination. **d)** Cre-/- and Cre+/- mice prefer to investigate the novel individual over a familiar individual (Two-way repeated-measures ANOVA Interaction Partner F(1,16)=27.49, p<0.0001; Šídák’s multiple comparisons test familiar vs. novel Cre-/- p=0.0030, Cre+/- p=0.0046). **c)** Cre-/- and Cre+/- mice show a significant preference to interact with the novel over the familiar individual (one-sample t-test against zero Cre-/- t=00695.818, df=9, p=0.0003; Cre+/- t=3.776, df=7, p=0.).

**Supplementary Figure 11.**
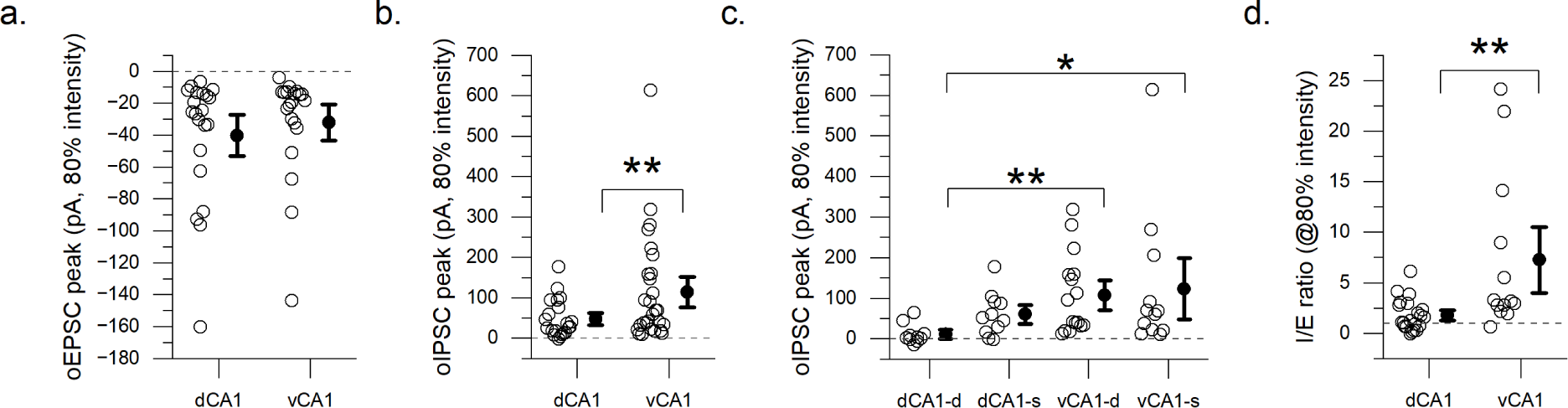
vCA2 demonstrates greater inhibition of vCA1 compared to dCA2 inhibition of dCA1. **a)** AVPR1b-Cre+/- subjects (dCA2: n = 3, vCA2: n=5) were injected with excitatory ChR2 to express specifically in dCA2 or vCA2. EPSC and IPSC responses were recorded in dCA1 or vCA1 under 80% intensity optic stimulation of dCA2 or vCA2, respectively. No difference was observed in EPSC peak in dCA2 projections to dCA1 compared to vCA2 projections to vCA1 (Mann-Whitney test, p=0.4614, dCA1: n =21, vCA1: n=20). **b)** A significantly higher IPSC peak was observed in vCA2 projections to vCA1 compared to projections to dCA2 projections to dCA1 (Mann-Whitney test, p=0.0036, dCA1: n =21, vCA1: n=28) **c**) The difference in inhibitory tone is apparent in vCA2 projects to both deep and superficial vCA1 layers (Kruskal-Wallis test, p=0.0028; Dunn’s multiple comparisons test, dCA1-d vs. dCA1-s p=0.0992, dCA1-d vs. vCA1-d p=0.0023, dCA1-d vs. vCA1-s p=0.0142, dCA1-s vs. vCA1-d p>1, dCA1-s vs. vCA1-s p>1, vCA1-d vs. vCA1-s p>1, dCA1-s: n=10, vCA1-d: n=11, dCA1-s: n=16, vCA1-s: n=12). **d**) There is a significantly higher inhibitory/excitatory ratio in projections from vCA2 to vCA1 compared to projections from dCA2 to dCA1 (Mann-Whitney test p=0.0031, dCA1: n =21, vCA1: n=13).

## Methods

**Table 1:** Software and Algorithms.

|  |  |  |
| --- | --- | --- |
| ANY-maze | Stoelting Co. | https://www.anymaze.co.uk/index.html |
| Prism | Graphpad | https://www.graphpad.com/scientific-software/prism/ |
| Fiji | ImageJ | https://imagej.net/software/fiji/ |
| Imaris | Oxford Instruments | https://imaris.oxinst.com/ |
| OriginLab | OriginLab Corporation | https://www.originlab.com/ |
| pClamp 10 | Molecular Devices | https://www.moleculardevices.com/products/axon-patch-clamp-system/acquisition-and-analysis-software/pclamp-software-suite |

### Immunofluorescent Labeling & Imaging

We perfused subject mice using saline followed by 4% PFA in ice-cold PBS. Brains were extracted and incubated in 4% PFA overnight. Brains were sliced in coronal, horizontal, or transverse orientation as specified with thickness of 60 µm using a Leica VT1000S vibratome. Post-hoc immunohistochemistry of in vitro electrophysiological experiments was performed by fixing 400 mm slices in 4% PFA at 4°C overnight. Sections were permeabilized and blocked for 1 hour with 5% goat serum and 0.4% Triton-X in PBS at room temperature. Sections were incubated overnight unless otherwise specified with the primary antibodies specified below at 4°C in 0.1% Triton-X in PBS plus 5% goat serum. The following day, slices were washed with PBS three times for 10 minutes in PBS and then incubated with secondary antibodies (all at concentration 1:500) for three hours. Slices were again washed three times in PBS for 10 minutes/wash. DAPI (ThermoFisher Scientific, #D1306) staining was applied at 1:1000 for 15 minutes in PBS at room temperature prior to mounting. Slices were mounted using Fluoromount (Sigma-Aldrich) and imaged using Zeiss LSM 700 confocal microscope.

### Avpr1bxAi14 labeling

Brains were dissected to extract the hippocampus from both hemispheres. The hippocampus was sliced in a transverse orientation. To get the complete series of expression, every third slice along the dorsoventral axis was selected for imaging. For quantification of prototypical CA2 marker expression, dorsal and ventral transverse slices were incubated with the following primary and secondary antibody combinations using the protocol described: PCP4 (1:300, rabbit anti-PCP4 #HPA005792, Sigma-Aldrich)/goat anti-rabbit IgG (Life Technologies) and either STEP (n-4, 1:1000, mouse anti-STEP # 4396, Cell Signaling Technology)/goat anti-mouse IgG1 (Life Technologies) or RGS14 (n=3, 1:500, mouse anti-RGS14 #75-170, Neuromab)/goat anti-mouse IgG2a (Life Technologies).

### PCP4-cre mouse line validation in vCA2

ThePCP4-cre mouse line (B6.Cg-Pcp4<tm2.1(cre)> (RBRC05662)) was developed by Toshitada Takemori and provided by RIKEN BRC through the National BioResource Project of the MEXT/AMED, Japan. Hippocampus were extracted from both hemispheres and sectioned in a transverse orientation (n=2). For the dual-labeling of Cre and PCP4, guinea pig anti-Cre (1:4000, Zuckerman Institute virus core) and rabbit anti-PCP4 (1:300, #HPA005792, Sigma-Aldrich) were used as the primary antibodies. Goat anti-guinea pig IgG-555 (1:500, # A-21435, Life Technologies) and goat anti-rabbit IgG-488 (1:500, # A-11008, Life Technologies) were used as the secondary antibodies.

### Cell counting

For basic quantification of CA2 markers in dorsal and ventral hippocampus, we chose transverse slices from the last 1/3 of the hippocampus (mouse n=5, slice n=7) and dorsal slices from the first 1/3 of the hippocampus (mouse n=7, slice n=8). The following coordinates were used to define CA2 across slices: from the tip of the mossy fiber, as visualized with PCP4 labeling of the mossy fibers, the lower edge of the area was defined as a 205 µm line lying along the lower edge of the pyramidal layer towards the DG. The deep-to-superficial edge of the area on the CA3-side was defined as 110 µm centered in the middle pyramidal layer. Finally, the upper edge of the area was defined by a 325 µm line moving towards CA1 from the defined CA3 border. 5 consecutive z-stack images with a delta of 2.04 µm were selected, for a final volume of 237,864 µm3/slice. These boundaries are consistent with a conservative estimate of CA2 transverse width in line with previous reports^25^.

### iDISCO brain clearing and light sheet microscope imaging

Avpr1b-Cre x Ai14 brains with heterozygous expression of Cre (n=3) were processed as previously described^49^. tdTomato expression was identified using the primary antibody rabbit anti-RFP (1:1000, Rockland, #600-401-379). Background labeling was captured with chicken anti-GFP (1:2000, AVES Labs, #GFP-1020) for 7 days followed by the following respective secondary antibodies: donkey anti-rabbit conjugated to Alexa-488 (1:1000, Thermo Fisher Scientific, # A-21206) and donkey anti-chicken conjugated to Alexa-647 (1:2000, Jackson #703-605-155) for 7 days. Light sheet microscopy was performed using an UltraMicroscope II light-sheet microscope (LaVision). 3-D reconstruction was completed via Imaris software (Bitplane).

### In-situ hybridization and RNAscope

#### Brain collection

Brains from Avpr1b-Cre+/- mice crossed to an Ai14 background to express tdTomato in Avpr1b-expressing cells were extracted (n=2) and immerse in dry-ice cold Butan X to freeze for 6 seconds. Frozen brain was embedded in OCT and sliced coronally in 16 µm sections using a Leica CM3050 S cryostat. Slices were mounted onto Superfrost Plus slides (12-550-15, FisherBrand). Sections were moved to a −20°C fridge to dry and then stored in an air tight wrapped slide box at −80°C.

#### Pre-assay tissue preparation

To prepare slices, slides were immersed in 10% NBS for 15 minutes at 4°C. Slides were then transferred to wash in different EtOH concentrations for 5 minutes each at room temperature (RT): 50%, 70%, 100%, 100%. Slices were removed and placed on absorbent paper to dry for 5 minutes. An ImmEdge hydrophobic barrier pen was used to draw a barrier around the slices. A HybEZ II Hybridization System oven (Advanced Cell Diagnostics) was set to −40°C. Dried slides were places on the HybEZ slide rack and 4 drops of protease IV was added per section and allowed to incubate at RT for 30 minutes. Liquid was removed and the slides were immediately places in a tissue tek slide rack in a tissue tek staining dish filled with 1x PBS, and this process was repeated 1x time.

#### Fluorescent RNAscope assay

Avpr1b and tdTomato probes (Advanced Cell Diagnostics RNAscope Probes Cat #480141 and 317041, respectively) were warmed to 40°C for 10 minutes and then cooled to RT. Probes were spun down and then mixed at a ratio of 1:50 (Avpr1b: tdTomato). Excess liquid was removed from slides and 4 drops of the probe were added to the slides. Slides were placed in the HybEZ oven tray and secured in the oven. They were incubated for 2 hours at 40°C. Slides were then removed, tapped to remove excess fluid, and immersed in 1x wash buffer for 2 minutes at RT twice. For each AMP, excess fluid was removed from the slide and then 4 drops were added proceeding from 1 to 4. Slides were placed in the HybEZ oven tray and incubated with the following times: 1: 30 minutes, 2: 15 minutes, 3: 30 minutes, 4: 15 minutes. Following AMP4, slides were immersed in 1x wash buffer for 2 minutes at RT twice. Excess fluid was removed by tapping slide and 4 drops DAPI was added to slides. Slides were incubated for 1 minute at room temperature. DAPI was removed and immediately Prolong Gold Antifade (w/o DAPI) was added to the slide and cover-slipped. Slides were stored in dark at 4°C until they were imaged using a Zeiss LSM 700 confocal microscope.

### Viral injection

Subject mice were given 5 mg/kg carprofen analgesic and anesthetized with 2-5% isoflurane. Subjects were placed in a stereotaxic frame and a craniotomy was drilled at the target coordinates. A glass pipette containing the virus was lowered to the desired depth. A Nano-Inject II (Drummond Scientific) was used to inject 23 nL virus at intervals of 10 seconds to obtain the desired volume. Following injection, five minutes were allowed to pass before the pipette was retracted. All viral injection coordinates are in mm with Bregma as reference.

### Dual-injection tracing experiment

Avpr1b-Cre+/- subject mice (n=4) were injected in dCA2 (AP −2.0, ML +/-1.8, DV −1.7) with 200nL of either: AAV9.CAG.FLEX.eGFP.WPRE.HGH (1×10^12 pp/mL, UPenn Vector Core, cohort 1, n=2) or AAV9.EF1a.dflox.hChR2(H134R).EYFP.WPRE.HGH (1×10^12 pp/mL, Addgene, cohort 2, n=2). Subject mice were additionally injected in vCA2 (AP −3.0, ML +/-3.2, DV −3.8) with 200nL of either: AAV2.EF1a.DIO.mCherry (1×10^12 pp/mL, UPenn Vector Core, cohort 1) or AAV9.EF1a.dflox.hChR2(H134R).mCherry.WPRE.HGH (1×10^12 pp/mL, Addgene, cohort 2).

We observed no difference in the projection patterns between cohorts. Cohort 1 was injected unilaterally in the right hemisphere; cohort 2 was injected bilaterally. Single-injection tracing experiment: Amigo2-Cre+/- subject mouse (n=1) was injected in vCA2 (AP −3.0, ML +/-3.2, DV - 3.8) with 200 nL AAV9.EF1a.dflox.hChR2(H134R).mCherry.WPRE.HGH (1×10^12 pp/mL, Addgene). For subjects in cohort 1 (n=2), brains were extracted and sliced in a coronal orientation. For cohort 2 (n=2), brains were extracted and divided into three parts by 2 slices: 1) a coronal slice at the intersection of the hippocampus and the lateral septum separating the brain into an anterior and posterior portion, and 2) a slice in the posterior portion separating the two hemispheres. The anterior portion was sliced in a horizontal orientation to observe projections to the lateral septum. The hippocampus from the right hemisphere was extracted and sliced in a transverse orientation. The posterior portion from the left hemisphere was sliced in a horizontal orientation. Through this method, we were able to visualize the projections from dorsal and ventral hippocampus both outside the hippocampus in coronal and horizontal sections, and within the hippocampus in coronal, horizontal, and transverse sections.

Viral tracing, single injection: The hippocampus was dissected from both hemispheres of the Amigo2-Cre mouse brain. The hippocampus was sliced in a transverse orientation and then labeled with PCP4 (1:300, rabbit anti-PCP4 #HPA005792, Sigma-Aldrich) as primary and goat anti-rabbit IgG (Life Technologies), for secondary.

### Pharmacogenetic silencing of ventral CA2

Avpr1b-Cre-/- and +/- or PCP4-Cre-/- and +/- subject mice were injected with a Cre-dependent virus expressing the inhibitory hM4Di designer receptor exclusively activated by designer drugs (iDREADD) [200nL bilaterally at 1.9×10^12 pp/mL with AAV2/8 hSyn.DIO.hM4D(Gi)-mCherry] bilaterally at coordinates for dorsal CA2 (AP - 2.0, ML +/-1.8, DV −1.7) or ventral CA2 (AP −3.0, ML +/-3.2, DV −3.8).

### Dorsal and ventral CA2 silencing

Three weeks post-injection, subject mice were habituated to saline intraperitoneal injections 4 days prior to testing, alternating the side of injection each day. One day prior to testing, subjects were weighed to ensure proper CNO dosage. CNO solution (Cayman Pharmaceuticals, CAS # 34233-69-7) was mixed on day of the experiment to reach a concentration of 1 mg/mL in sterile saline. 30-minutes prior to each test, intraperitoneal injections of subject mice were performed with a volume to ensure 10mg/kg CNO dose unless otherwise specified. Subjects and any stimuli mice were moved to the testing area 30 minutes prior to the experiment start to habituate to the test room. Stimulus mice were always sex- and age-matched. Videos of experiments were recorded using a FireWire camera (DMK 31AF03-Z2; The Imaging Source) triggered through ANY-maze software.

### Behavior

Mice were housed with littermates unless otherwise noted, and kept on a 12-hour light-dark cycle in air-filtered, temperature- and humidity-controlled conditions with food and water available ad libitum. Behavior was run under low red-light conditions unless otherwise specified. All behavior experiments were recorded with ANY-maze software.

### Social recall test

One day prior to testing, subjects were habituated to the arena, a three-chambered box with each chamber measuring 19 cm x 40.5 cm x 22 cm. The apparatus was made of clear Plexiglass with openings 10 cm side to allow access across chambers. Within the arena, two empty pencil cups (radius 5 cm) were placed in each of the side chambers.

Following 10 minutes habituation, subjects were returned to their home cages. On the test day, following CNO injection, subject mice and littermates were separated into two testing cages at random (A or B) and brought to the test room. Stimulus mice, with whom the subject mice had never interacted, were also brought to the test room to habituate. After 30-minutes, the subject mice were introduced to the central chamber of the three-chamber apparatus with two barriers blocking the doorways to the side chambers, which each held an empty pencil cup. The barriers were removed to start the test, and the subject mice freely explored the three chambers for 10 minutes. The mouse was again confined to the central chamber. A littermate from the opposite testing cage was placed under the pencil cup in one chamber, and a novel stimulus mouse was placed under the pencil cup in the opposite chamber. The location of the novel and littermate mouse was alternated across mice. The barriers were again removed, and the subject allowed to explore the novel conspecific and familiar littermate freely for 10 minutes. The time spent in each chamber was measured automatically using zones defined within ANY-maze.

### Social discrimination test

One day prior to testing, mice were habituated to an oval arena that consisted of two half-circles with radius 15cm connected to a central square area with length of 30 cm (dimensions: length 60 cm, width 30 cm, height 45 cm). Wire pencil cups (radius 5 cm) were placed 10 cm from the two ends of the arena along the midline. Following 10-minute habituation, subjects were returned to their home cages. The next day following CNO injection and 30-minute habituation to the testing room, subjects were re-introduced to the oval arena with empty pencil cups for 5 minutes. The subject was removed and returned to their home cage, and two stimulus mice were placed into the arena, with one under each pencil cup. The subject mouse was returned to the arena and allowed to freely interact with these individuals for 5 minutes. The subject mice were then returned to their home cage for 30 minutes. The subject was then returned to the oval arena with the empty pencil cups for 5 minutes. The subject was again removed to their home cage, and two stimulus mice (one from the prior trial and a third novel individual) were placed at random under each of the two pencil cups. The subject mouse was again returned to the arena and allowed to investigate the now-familiar and novel individual freely for 5 minutes. The time in a circular zone (radius = 10cm) around each cup was automatically scored.

### Avpr1b-Cre and PCP4-Cre Behavior validation

For dCA2 pharmacogenetic silencing, Avpr1b-Cre mouse brains were sliced coronally to allow all dorsal hippocampal subfields to be simultaneously imaged. For vCA2 pharmacogenetic silencing, Avpr1b-Cre or PCP4-Cre mouse brains were sliced horizontally to allow all ventral hippocampal subfields to be simultaneously imaged. Brains were incubated with the following primary/secondary antibody pairing: PCP4 (1:300, rabbit anti-PCP4 #HPA005792, Sigma-Aldrich)/goat anti-rabbit IgG (Life Technologies). Mice with low levels of expression (<50% Avpr1b-expressing CA2 pyramidal neurons infected in Avpr1b-Cre mice or <50% infection of DAPI+ cells in vCA2 region in PCP4-Cre mice) or significant expression outside of CA2 (>10% florescent expression of the areas of CA1, CA3 or DG with a binary mask in PCP4-Cre mice). Due to the absence of an Avpr1b antibody to confirm co-expression with iDREADD, we used PCP4 to label vCA2. As noted in the text, we expect two major populations of vCA2 neurons: PCP4+/Avpr1b+ (59.8 ± 7.5%) and PCP4+-only (36.6 ± 6.0%). Thus, we expect roughly 59.8% of pyramidal neurons infected in the Avpr1b-Cre mouse to express the iDREADD virus. Therefore, mice with <29.9% expression of iDREADD (50% of the Avpr1b-expressing population) in vCA2 from either hemisphere were excluded from analysis in experiments in which the Avpr1b-Cre subjects were used.

### Social aggression

PCP4-Cre+/- male mice and their Cre-/- littermates were bilaterally injected with AAV8-hm4Di-mCherry into the vCA2. After 8 days of recovery, the mice were singly housed for 10 days before the social aggression test. The resident-intruder paradigm was adjusted based on previously published experiments^40^. Stimulus mice (7-8 week-old BALB/cByJ male intruders) were group housed and used only once per day. On the aggression screening days, the subject mice were first received a saline injection (5 ml/kg, ip) as a control for the test day. After a 30-min room habituation, stimulus mice were introduced to the home cages of the subject mice for 5-min free interaction. In accordance with Columbia IACUC rules, the attack was allowed to continue for 1 min after its onset, which was defined by a clear bite. To increase the occurrences of aggression and select the aggressive subjects, 3 continuous days of screenings were allowed for each subject (19-21 days after the virus injections). On the test day, CNO (5 mg/kg, ip) was injected 30 min prior to the test. A 10-min resident-intruder test was then performed with a novel intruder introduced to the home cage of the PCP4-Cre subject mice. ANY-maze software (Stoelting Company) was used for recording and measurements. PRISM (Graphpad) was used for data analysis.

### In vitro electrophysiology and data analysis

*In vitro* electrophysiology was performed in the 8-16 week-old Avpr1b-Cre x Ai14 mice and Avpr1b-Cre mice. Mice were anesthetized by isoflurane inhalation and transcardialy perfused with ice-cold sucrose ACSF containing, in mM: 195 sucrose, 10 glucose, 25 NaHCO3, 2.5 KCl, 1.25 NaH2PO4, 2 sodium pyruvate, 0.5 CaCl2, 7 MgCl2. Mice were then decapitated and hippocampi were quickly dissected in the sucrose ACSF, placed into a 4% agar mold, and sectioned into 400 µm transverse slices with a vibratome (VT1200S, Leica). Slices were then transferred into a warmed (32°C) recovery beaker containing 50% sucrose ACSF and 50% ACSF (2.5 glucose, 125 NaCl, 25 NaHCO3, 2.5 KCl, 1.25 NaH2PO4, 3 sodium pyruvate, 1 ascorbic acid, 2 CaCl2, and 1 MgCl2) for recovery. After at least 30 min of incubation, the beaker was kept in room temperature for an additional 30 min prior to recording. All the solutions were continuously saturated with carbogen gas (95% O2/ 5% CO2).

Recording pipettes and stimulation pipettes were prepared with borosilicate glass capillaries using a heated-filament puller with an open tip resistance of 3-6 mOhm when filled with an intracellular solution containing, in mM: 135 potassium gluconate, 5 KCl, 0.2 EGTA, 10 HEPES, 2 NaCl, 5 MgATP, 0.4 Na2GTP, 10 Na2phosphocreatine, and biocytin (0.2% by weight), adjusted to pH 7.2 and osmolarity 295 mOsm. For voltage clamp recordings of in vitro optogenetics, KCl was replaced with cesium methanesulfonate in the intracellular solution. Whole cell recordings were acquired using a Multiclamp 700A amplifier (Molecular Device), data acquisition interface ITC-18 (Instrutech) and pClamp 10. CA1/CA2 border was identified as the end of the stratum lucidum (SL) and the edge of Cre-dependent expression of fluorescent. Pyramidal neurons were identified based on the cellular morphology and intrinsic properties such as the absence of fast spiking. Access resistance (15-25 MΩ) was continuously monitored.

Membrane capacitance was obtained at −70mV from a V-shape voltage clamp ramp protocol with ΔV=±10mV. Input resistance was calculated from a voltage step from −70mV to - 80mV in voltage clamp configuration. Voltage sag was measured with the negative current step initiated from −70mV in which the steady hyperpolarization potential was between −97mV and - 103mV. Sag ratio was calculated as (V_peak_-V_stt_)/(V_peak_-70), where V_peak_ is the minimum potential at the beginning of the step and V_stt_ is the steady hyperpolarization potential at the end of the step. Rheobase current and action potential properties were calculated with a 1000pA, one-second current ramp initiating from −70mV. For the experiments involving optogenetic activation of CA2 neurons, Avpr1b-Cre+/- mice were injected with either a AAV8-EF1a-DIO-hChR2(H134R)-mCherry (Addgene, #20297, 200nL bilaterally at 10^12 pp/mL) targeting ventral CA2 (AP −3.0, ML +/-3.2, DV −3.8), or a AAV5-EF1a-DIO-hChR2(H134R)-eYFP (Addgene, #20298, 200nL bilaterally at 10^12 pp/mL) targeting dorsal CA2 (AP −2.0, ML +/-1.8, DV −1.7) three weeks prior to the experiments. During the experiments, photostimulation was delivered as a 2-ms pulse of 470 nm light with a LED light source (ThorLabs High-power 5-channel LED driver with pulse modulation, model DC2100) through a 40x immersive objective with the recorded cell centered in the illumination field. For voltage clamp recording, cells were clamped at −75 mV (for EPSCs), −70 mV (for contingency of success rate), and +5 mV (for IPSCs). For current clamp recording, cells were held at −70 mV. Electrical stimulation was generated as a 2-ms pulse. The following drugs were bath-applied at the following concentrations: SR 95531 (1 µM, Tocris, #1262), CGP 55845 (2 µM,Tocris, #1248). Data analysis was performed using custom software written in OriginC^50^ (OriginLab Corporation, Northampton, MA) and PRISM (Graphpad)^50^.

To verify the location and the morphology of recorded neurons, all acute hippocampal slices were fixed, permeabilized and blocked for biocytin labeling at the conclusion of recordings. Biocytin was visualized by incubating hippocampal slices with Alexa Fluor 647-conjugated streptavidin (1:1000, Invitrogen#S21374).

